# Fes-deficient macrophages enhance CD8^+^ T cell priming and tumour control through increased pro-inflammatory cytokine production and localization

**DOI:** 10.1101/2024.02.27.581601

**Authors:** Brian J. Laight, Danielle Harper, Natasha Dmytryk, Shengnan Zhang, Andrew Garven, Changnian Shi, Richard Nauman, Jacob Kment, Faizah Alotaibi, Ivan Shapavalov, Yan Gao, Jeff Mewburn, Caitlyn Vlasschaert, David LeBrun, Kathrin Tyryshkin, David Berman, Amber Simpson, Charles Graham, Andrew W. Craig, Sameh Basta, Madhuri Koti, Peter A. Greer

## Abstract

Homeostatic immunoregulatory mechanisms that prevent adverse effects of immune overaction can serve as barriers to successful anti-cancer immunity, representing attractive targets to improve cancer immunotherapy. Here, we demonstrate a novel role of the Fes tyrosine kinase, abundantly expressed in immune cells, as an innate intracellular immune checkpoint. Host Fes-deficiency delays tumour onset in a gene dose-dependent manner and improves murine triple negative breast cancer and melanoma tumour control, survival, doxorubicin efficacy, and anti-PD-1 therapy sensitization. These effects were associated with a shift to an anti-tumourigenic tumour immune microenvironment. *In vitro*, we observed increased Toll-like receptor signaling, and proinflammatory cytokine production and presentation from antigen presenting cells, leading to increased T cell activation, cancer cell killing and tumour control. This study highlights Fes as a novel innate immune checkpoint with potential as a predictive biomarker for effective immune checkpoint blockade treatment, and a potential therapeutic target to improve anti-cancer immunotherapy.

## Introduction

Immune checkpoint inhibitors (ICI)^1,2^ which have primarily been focussed on enhancing adaptive cytotoxic T lymphocyte (CTL) activity through inhibiting the immune checkpoints CTLA-4 and PD-1 on the surface of T cells have revolutionized patient care. Unfortunately, ICIs are not a panacea; and the heterogeneity of tumours and of infiltrating immune cell types (some immunosuppressive) provides both barriers to successful cancer immunotherapy^3^ and opportunities for improvement. Strategies to overcome the immune resistance exhibited by many tumour types include searching for novel immune checkpoints^4^ or targeting different cell types within the tumour microenvironment (TME), including other immune cells^5,6^.

Initiation of adaptive anti-cancer immune responses involves antigen presenting cells (APCs), which engulf dying cancer cells and subsequently recruit and prime CTLs. Many commonly used therapies, such as anthracyclines (e.g., doxorubicin [Dox]) or radiotherapy^7^, can initiate such responses through immunogenic cell death (ICD). ICD induces the release of tumour-associated antigens and damage associated molecular patterns (DAMPs) from dying cancer cells, activating APCs and allowing them to prime an adaptive anti-cancer immune response. For instance, high mobility group box 1 (HMGB1) protein released from dying cancer cells during ICD binds APC Toll-like receptor 4 (TLR4) and activates signaling cascades required for pro-inflammatory cytokines production^8–10^. These pro-inflammatory cytokines, colloquially known as Signal 3, play a crucial role in CTL priming; and their absence is associated with deficient CTL effector functions and reduced memory formation^11,12^. Additionally, some APCs (e.g., macrophages) have been shown to play important roles in direct tumour cell killing^13,14^. Enhancing inflammatory signaling to activate the anti-cancer functions of APCs is an attractive strategy for enhancing adaptive anti-cancer immunity^14,15^.

While pro-inflammatory cytokine production is crucial for priming anti-cancer immune responses, uncontrolled release of these cytokines can have detrimental and even fatal consequences; such as septic shock^16^, auto-immune disorder^17^, or cytokine storm^18^. To combat these negative consequences, various mechanisms have evolved to attenuate immune responses. The non-receptor tyrosine kinase Fes appears to play such a role, and is abundantly expressed in APCs^19^. Fes limits TLR4 signaling^20^, in part through facilitating TLR4 internalization, preventing sustained stimulation^21^. Fes has also been shown to associate with MHCI and suppresses inflammatory signaling in macrophages through activation of the SH2 containing protein tyrosine phosphatase-2 (SHP2)^22^. While mechanisms to suppress immune cell activation are important to maintain homeostatic conditions^23^, in cancer contexts these mechanisms may enable tumour growth and limit the efficacy of immunostimulatory therapeutics^24^.

Following previous observations implicating Fes in regulation of TLR internalization^21^, involvement in vesicle trafficking^25^, and signalling suppression through the MHC-I axis^22^, we sought to explore its involvement in regulating anti-cancer immune responses. In this study, we identify Fes as an innate intracellular immune checkpoint. We show that Fes-deficiency is associated with delayed tumour initiation, correlating with increased immune activity within the premalignant niche. Fes-deficiency in mouse syngeneic triple negative breast cancer (TNBC) and melanoma models improved tumour control and survival, which was further enhanced by Dox treatment. Tumours in Fes-deficient mice displayed increased activated and PD-1 positive CTLs, which correlated with sensitization of tumours to ICI therapy. We attributed this, in part, to increased T cell priming capability of *fes^-/-^*bone-marrow derived macrophages (BMDMs) compared to *fes^+/+^*, due in part to not only an increase in Signal 3 cytokines production, but also the method of interleukin-12 (IL-12) presentation to CTLs through macrophage membrane retention. Taken together, these results demonstrate a novel role for Fes as an immune checkpoint, and a potential predictive biomarker and target for improved ICI therapy.

## Results

### Fes-deficiency delays tumour growth in a gene dose-dependent manner and is associated with increased immune infiltration in a transgenic mouse model of Her2^+^ breast cancer

While Fes-deficiency in an immunodeficient mouse engraftment model, showed delayed tumour growth^26^, it did not provide insight into its role in adaptive immunity. Therefore, we assessed HER2^+^ tumourigenesis in an *MMTV-Neu* transgenic mouse model^27^ crossed with transgenic mice harbouring a targeted kinase-inactivating K588R knock-in mutation in *fes* (*fes^KR^)*^28^. The average tumour onset time in *fes^+/+^* mice was 274 days and was delayed to 316 days in *fes^KR/+^* and 389 days in *fes^KR/KR^* mice (Fig.1a) demonstrating that a functional inhibition of Fes could delay tumour onset. Interesting, once palpable tumours were detected, their growth rates were indistinguishable (Extended Data Fig.1a). This Fes-associated delay in tumourigenesis in mice correlates with higher human cancer incidence rates in the UK Biobank cohort (HR 1.03 [1.01 – 1.05], p=0.01) and risk of death (HR 1.06 [1.01 – 1.11], p=0.03) in subjects homozygous for a single nucleotide polymorphism associated with a 50% reduction in *FES* expression (rs17514846)^29^.

**Fig. 1:**
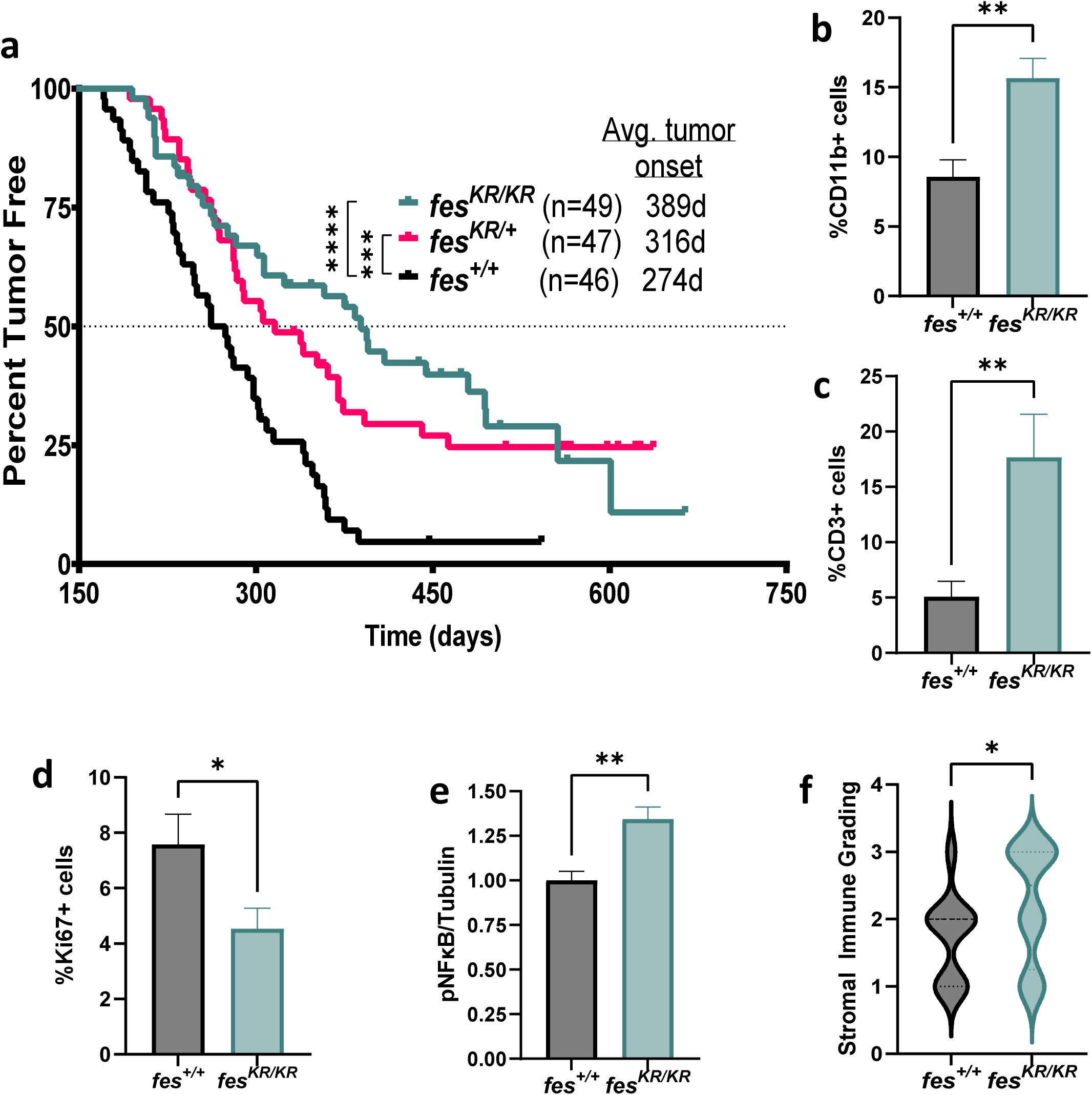
Fes-deficiency delays tumour growth and is associated with increased immune infiltration in a transgenic mouse model of Her2^+^ breast cancer. **a,** *MMTV-Neu* transgenic mice wild type for *fes* (*fes^+/+^*, n=46), or harbouring one (*fes^KR/+^*, n=47) or two (*fes^KR/KR^*, n=49) kinase-dead K588R alleles, were monitored for tumor onset by mammary gland palpation. **b-d**, pre-malignant mammary gland sections were assessed by immunohistochemistry and quantification of % cells staining for: **b,** CD11b (*fes^+/+^*, n=3; or *fes^KR/KR^*, n=3); **c,** CD3 (*fes^+/+^*, n=6; or *fes^KR/KR^*, n=6); or **d,** Ki67 (*fes^+/+^*, n=3; or *fes^KR/KR^*, n=3). **e,** pre-malignant glands were isolated and probed for pNFκB by western blot analysis (*fes^+/+^*, n=5; or *fes^KR/KR^*, n=5). **f,** mammary gland sections of similarly sized tumours were stained with H&E and evaluated by a blinded pathologist to assess stromal immune grading (*fes^+/+^*, n=11; or *fes^KR/KR^,* n=24). Time-to-event (**a**) statistics were assessed through Log-rank (Mantel-Cox) test. IHC (**b-d**) and western blot (**e**) statistical analysis was performed through an unpaired one-tailed t-test. *p<0.05, **p<0.01, ***p<0.001, ****p<0.0001.

To explore a potential role of the immune system on tumour onset delay in the transgenic mouse model, we isolated pre-malignant mammary glands and performed immunohistochemical staining for CD11b, CD3, and Ki67. This revealed an increase in both CD11b+ (Fig.1b) and CD3+ cells (Fig.1c), and a decrease in Ki67+ cell in *fes^KR/KR^* glands (Fig.1d); demonstrating increased immune infiltration and reduced mammary epithelial cell proliferation. Immunoblotting analysis revealed an increase in phosphorylated NFκB in *fes^KR/KR^* mammary glands (Fig.1e); a transcription factor which promotes expression of inflammatory cytokines. Finally, an increase in immune cell infiltration was also apparent after tumours formed, where there was an increase in immune cell content in *fes^KR/KR^* mice (Fig.1f).

### Fes-deficient mice have delayed tumour growth, improved response to doxorubicin treatment and prolonged survival

Reduced functional Fes was associated with delayed tumour onset and an increase in CD11b and CD3-infiltrating cells, and NFκB phosphorylation, implicating Fes in the regulation of anti-tumour immunity. To further explore this, we employed E0771 mammary carcinoma and B16F10 melanoma syngeneic orthotopic engraftment models using C57BL/6 mice targeted with a *fes* null mutation (*fes^-/-^*) or wild type (*fes^+/+^*) mice. Dox treatment served to induce ICD and promote innate and adaptive anti-tumour immunity (Fig.2a,d). Fes-deficiency was associated with improved tumour control (Fig.2b,e) and survival (Fig.2c,f) in the absence of treatment, and was further enhanced with Dox treatment in both models. Dox treatment prolonged survival of *fes^-/-^* mice in both models. However, in *fes^+/+^* mice, Dox treatment only improved survival in the B16F10 model. Since the same cancer cells are used in engraftments of both *fes^-/-^* or *fes^+/+^*mice, the observed differences are consistent with a Fes effect driven by the tumour microenvironment. Furthermore, Fes expression is very low in E0771 cells and is undetected in B16F10 cells (Extended Data Fig.2b).

**Fig. 2:**
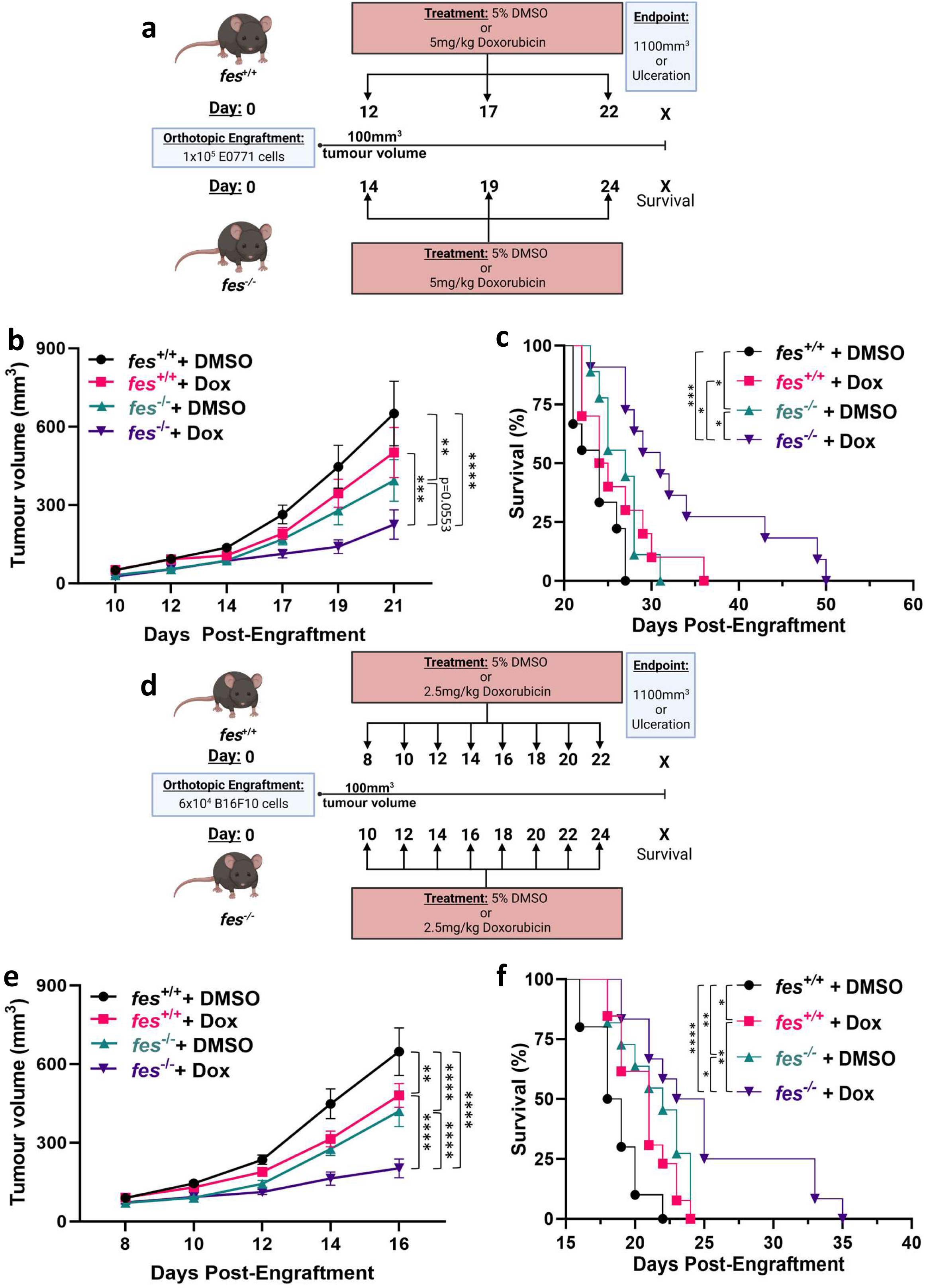
Fes-deficient mice have delayed engrafted tumour growth, improved response to doxorubicin treatment and prolonged survival. **a,d,** Schematics of tumour growth (**b,e**) and survival analysis (**c,f**) of E0771 (**a-c**) and B16F10 (**d-f**) syngeneic engraftment models, respectively. Cancer cells were orthotopically engrafted into either *fes^+/+^* or *fes^-/-^* mice (female for E0771 and male for B16F10) and treated with either 5% DMSO (vehicle) or Dox (E0771, 5mg/kg; B16F10, 2.5mg/kg). (**b,e**) Tumor growth was monitored using calipers to an endpoint of 1.1cm^3^ or physiological endpoint (e.g., ulceration), and (**c,f**) survival was plotted. Data are represented as tumour volumes ± SEM of two independent experiments of both E0771 and B16F10 models with the following numbers of animals per group: *fes^+/+^* + DMSO (E0771, n=9; B16F10, n=10), *fes^+/+^* + Dox (E0771, n=10; B16F10, n=13), *fes^-/-^* + DMSO (E0771, n=9; B16F10, n=11), *fes^-/-^*+ Dox (E0771, n=11; B16F10, n=12). Tumour growth results (**b,e**) were analyzed by two-way ANOVA with Tukey’s multiple comparisons test and survival data (**c,f**) was analyzed using Log-rank (Mantel-Cox) test. * p<0.05, ** p<0.01, *** p<0.001, ****p<0.0001.

### Fes-deficiency correlates with a more activated anti-tumoural immune microenvironment, which is enhanced by doxorubicin therapy

The previous observations implicated Fes in regulating an anti-cancer immune response, which prompted comparisons of the tumour immune profiles in *fes^+/+^* and *fes^-/-^* mice in these E0771 and B16F10 engraftment models (Fig.3a,h). There was no genotype associated differences in the frequency of tumour-associated CD8^+^ T cells (CTLs) (Fig.3b,i), or splenic CTLs (Extended Data Fig.2c,m) in either engraftment model. However, tumour-associated CTLs displayed higher frequencies of activation (CD69^+^) in *fes^-/-^* mice, and this phenotype was further elevated with Dox treatment (Fig.3c,j). *fes*^-/-^ mice also had higher levels of splenic CTL activation, but this was not enhanced by Dox treatment (Extended Data Fig.2d,n). Increased tumour-associated CTL PD-1 positivity (indicative of more extensive and prolonged activation) was also seen in *fes^-/-^* mice with Dox treatment in both models (Fig.3d,k). PD-1 positivity was inducible by Dox treatment in E0771 tumours only in *fes^-/-^* mice (Fig.3d). Interestingly, PD-1 positivity without Dox treatment was already higher in *fes^-/-^* mice in the B16F10 model, and this was not further induced by Dox treatment (Fig.3k). In spleens, there was a Dox-induced increase in PD-1 positivity only in *fes^+/+^* treated animals with the E0771 model (Extended Data Fig.3e,o).

**Fig. 3:**
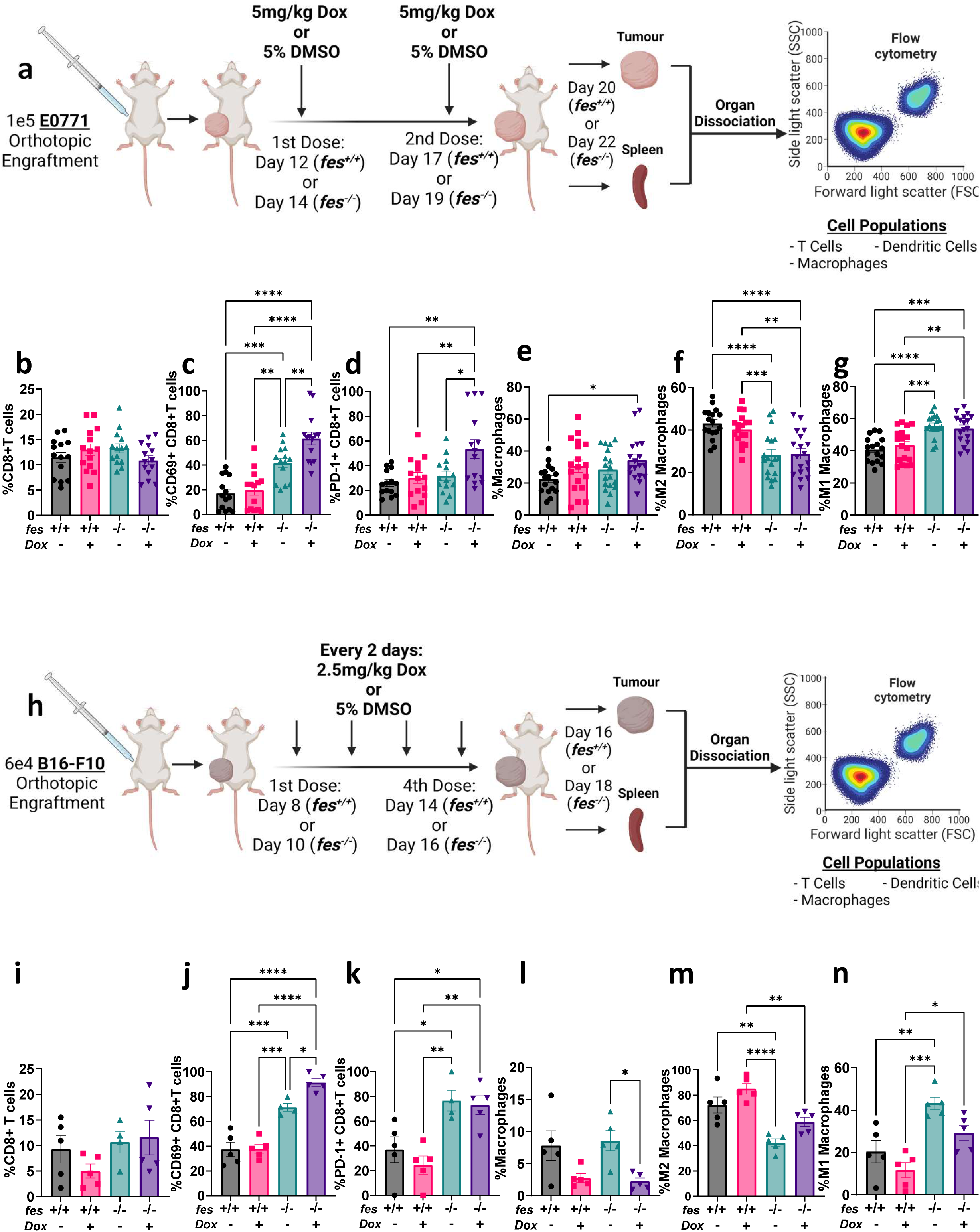
Fes deficiency promotes a shift to an anti-tumour microenvironment and sensitizes tumours to doxorubicin-induced T cell activation. **a,h,** Schematic representations of immune profiling of E0771 or B16F10 tumour models, respectively. Tumours were dissociated and analyzed by flow cytometry for: **b,i,** total CD8+ T cells (CD45+, CD3+, CD8+;[CTLs])(E0771, n=14; B16F10, n=5); **c,j,** CD69+ CTLs (E0771, n=14; B16F10, n=5); **d,k,** PD-1+ CTLs (E0771, n=14; B16F10 n=5); **e,l,** total macrophages (CD45+, CD11b+, F4/80+)(E0771, n=18; B16F10 n=5); **f,m,** M2 macrophages (CD206+)(E0771, n=18; B16F10, n=5); and **g,n,** M1 macrophages (CD206-, MHCII+ and/or CD80+)(E0771, n=18; B16F10, n=5). Results were analyzed by one-way ANOVA with Tukey’s multiple comparisons test; * p<0.05, ** p<0.01, *** p<0.001, **** p<0.0001.

We next looked at the frequency of tumour-associated macrophages and saw no *fes* genotype associated differences in either model (Fig.3e,l). However, we observed a *fes* genotype dependent decrease in immunosuppressive M2 (CD206^+^) macrophages (Fig.3f,m) and a subsequent increase in proinflammatory M1 (CD206^-^/CD80^+^ and/or MHCII^+^) macrophages (Fig.3g,n) in *fes^-/-^* mice in both models. This shift was also seen in the spleen (Extended Data Fig.3g,h,q,r). In both tumor models, we saw no *fes* genotype associated changes in dendritic cell (DC) numbers (Extended Data Fig.3k,u). However, in the E0771 model, we observed increased DC activation in tumours in *fes^-/-^* mice (Extended Data Fig.3l), but this was not observed in the B16F10 model (Extended Data Fig.3v). These results demonstrate that Fes-deficiency is associated with a more anti-tumourigenic immune profile in both E0771 and B16F10 models.

### Tumours in Fes deficient mice are more sensitive to PD-1 therapy

In light of our observation of increased PD-1 positivity on tumour-associated CTLs in *fes^-/-^* mice with both engraftment models (Fig.3d,k), we decided to explore the effect of anti-PD-1 therapy. Since CTLs in the E0771 model only showed an increase in PD-1 positivity following Dox therapy, similar to previous studies^30^, we treated all mice with Dox (to induce ICD), and either an isotype antibody control (IgG) or anti-PD-1 antibody (Fig.4a,d). In both the E0771 and B16F10 models, anti-PD-1 treated *fes^+/+^* mice showed improvements in tumour control (Fig. 4b,e), but not in survival (Fig.4c,f) compared to IgG. This observation is consistent with literature describing these tumour types as anti-PD-1 therapy resistant^31–36^. Impressively, Dox plus IgG treatment of *fes^-/-^* mice had the same (Fig.4b) or improved (Fig.4e) tumour control than Dox plus anti-PD-1 treatment of *fes^+/+^*mice. While anti-PD-1 therapy only marginally delayed tumour growth in *fes^+/+^*mice, anti-PD-1-treated *fes^-/-^* mice demonstrated greater tumour delay and a greater survival advantage compared to all other treatment groups (Fig.4b,c,e,f). These observations showed that Fes-deficiency sensitizes these resistant tumours to ICI therapy.

**Fig. 4:**
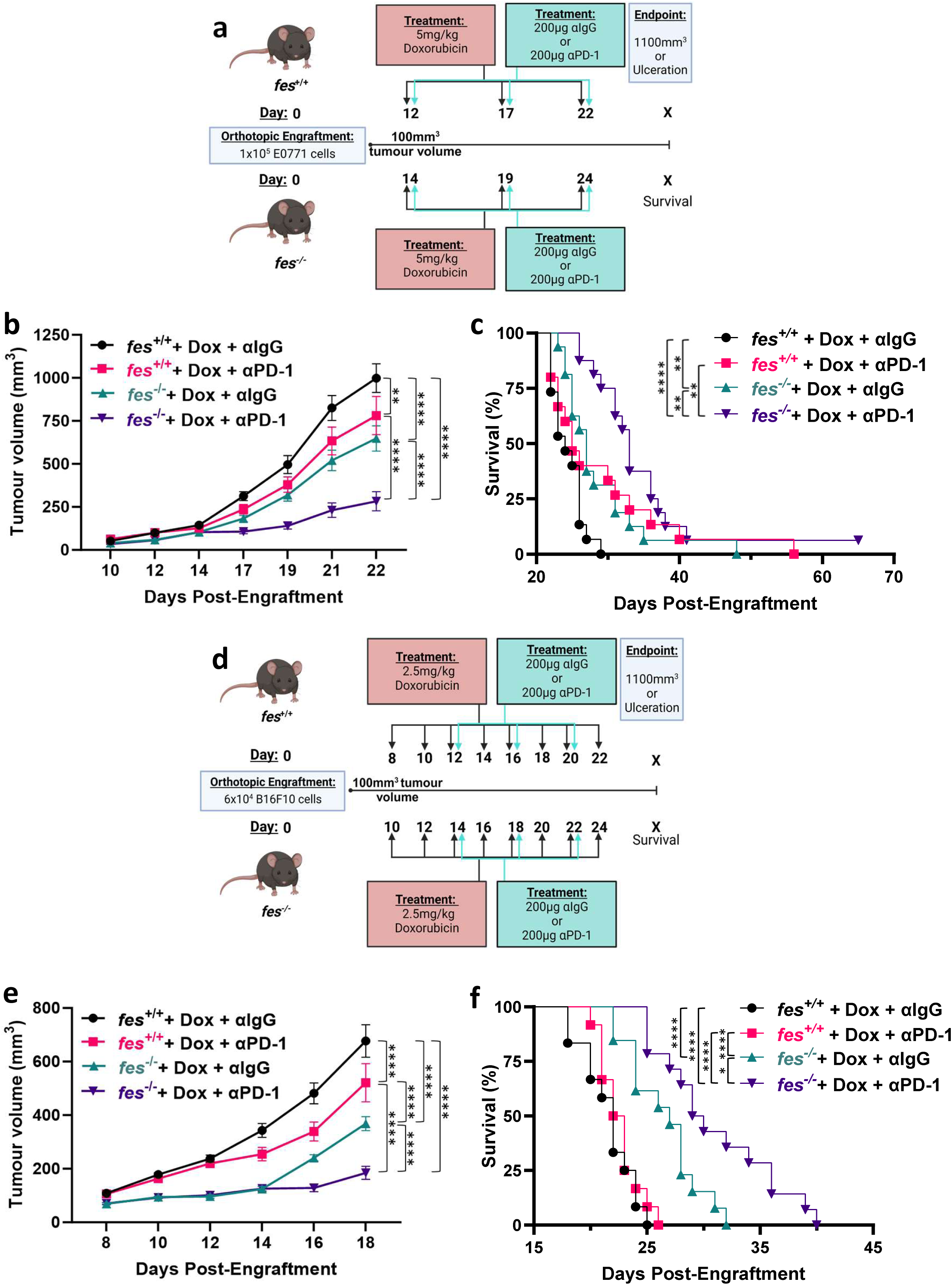
Fes deficiency enhances anti-PD-1 therapy and sensitizes resistant tumours. **a,d,** Schematics of anti-PD-1 treatment in the E0771 and B16F10 engraftment models, respectively. Cancer cells were orthotopically engrafted into either *fes^+/+^* or *fes^-/-^* mice (female for E0771 and male for B16F10). All mice were treated with either 5mg/kg Dox (E0771 model) or 2.5mg/kg Dox (B16F10 model), and treated with either 200µg IgG control antibody, or 200µg anti-PD-1 antibody. (**b,e**) Tumour growth was monitored using calipers to an endpoint of 1.1cm^3^ or humane endpoint (e.g., ulceration); and (**c,f**) survival was plotted. Data were represented as tumours volumes ± SEM of two (B16F10) or three (E0771) independent experiments with the following numbers of animals per group, respectively: *fes^+/+^* + Dox + αIgG (E0771, n=15; B16F10, n=12); *fes^+/+^* + Dox + αPD-1 (E0771, n=15; B16F10 n=12); *fes^-/-^* + Dox + αIgG (E0771, n=16; B16F10, n=13); *fes^-/-^* + Dox + αPD-1 (E0771, n=16; B16F10, n=14). Tumour growth (**b,e**) results were analyzed by two-way ANOVA with Tukey’s multiple comparisons test and survival data (**c,f**) was analyzed using Gehan-Breslow-Wilcoxon Test. * p<0.05, ** p<0.01, *** p<0.001, **** p<0.0001.

### Fes deficient macrophages have increased TLR signaling, increased pro-inflammatory cytokine production and altered IL-12 localization

Fes displays tissue-specific expression, with highest levels observed in immune cells, primarily macrophages and DCs, as well as NK and B cells; but Fes is not expressed in T cells, and to very low levels in E0771 cells (Extended Data Fig.2b). The host *fes* genotype associated changes in the phenotype of tumour-associated macrophages and enhanced CTL activation in these tumors, lead us to next explore Fes functions in macrophages, given their potential role as APCs.

We have previously shown that Fes plays a role in downregulating inflammatory signaling in macrophages, in part through promoting LPS-induced endocytosis of the TLR4 receptor^20^. Additionally, it has been proposed that Fes participates in suppressing TLR signaling through association with MHC-I and phosphorylation of SHP2; leading to suppression of inflammatory cytokine expression^22^. Here we explored the effects of Fes deficiency on the LPS-TLR4 signaling pathway in macrophages by culturing BMDMs from *fes^+/+^* and *fes^-/-^* mice and looked at downstream signaling by immunoblotting analysis following LPS stimulation. *fes^-/-^* BMDMs displayed earlier and more robust phosphorylation of NFκB (Fig.5a,b), TBK1 (Fig.5a,c), and IRF3 (Fig.5a,d) compared to *fes^+/+^* BMDMs. *fes^-/-^*BMDMs are also more sensitive to LPS than *fes^+/+^* BMDMs, with increased NFκB and TBK1 phosphorylation at concentration as low as 300pg/mL (Extended Data Fig.4a-c).

**Fig. 5:**
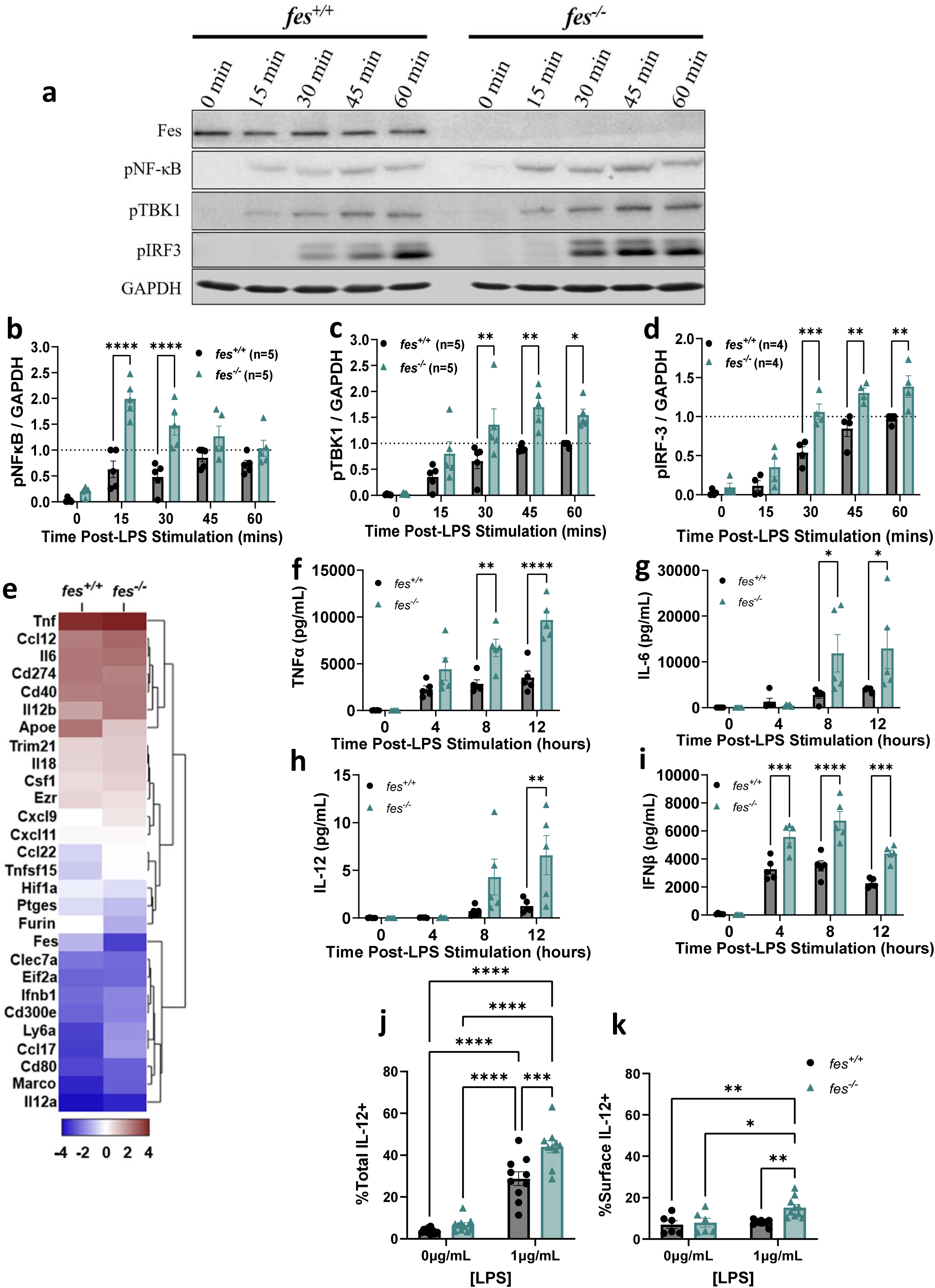
Fes deficient macrophages have increased TLR signaling, increased pro-inflammatory cytokine production and altered IL-12 localization. **a,** BMDMs from either *fes^+/+^* or *fes^-/-^* mice were stimulated with 1µg/mL LPS for the indicated time points before lysis and immunoblotting analysis of Fes, pNFκB (n=5), pTBK1 (n=5), pIRF3 (n=4), and GAPDH. Densitometry was assessed using ImageJ, and results were normalized to the band with the greatest density in the *fes^+/+^* BMDMs for either; **(b)** pNFκB, **(c)** pTBK1, **(d)** pIRF3. **f,** Bulk RNAseq analysis was performed on *fes^+/+^* (n=3) or *fes^-/-^* (n=3) BMDMs that were either untreated or treated with 1µg/mL of LPS for 4 hours. Selected transcripts with significantly different LPS-inducible levels in *fes^+/+^* and *fes^-/-^* BMDMs are illustrated. **g-j**, Secreted **(g)** TNFα, **(h)** IL-6, **(i)** IL-12, and **(j)** IFNβ cytokine levels were measured by LEGENDplex analysis conditioned media from *fes^+/+^* and *fes^-/-^* BMDMs (n=5 for each) untreated or stimulated with 1µg/mL LPS for 4, 8 or 12hrs. **k,l,** IL-12 expression and localization in *fes^+/+^*(n=10) or *fes^-/-^* (n=10) BMDMs unstimulated or stimulated with 1µg/mL LPS for 24 hours was assessed by flow cytometry. **(k)** Total levels were assessed by staining after permeabilization. **(l)** Cell surface levels were assessed by staining prior to permeabilization. Data in panels **b-e** and **g-l** were analyzed by two-way ANOVA with Sidak’s multiple comparisons test. * p<0.05, ** p<0.01, *** p<0.001, **** p<0.0001.

Enhanced LPS-induced phosphorylation and activation of NF-κB, TBK1, and IRF3, which play key roles in transcription of inflammatory cytokine genes, led us to analyze the transcriptome by bulk RNAseq from *fes^+/+^* or *fes^-/-^* BMDMs before or after a 4 hour stimulation with LPS. Increased LPS-induced RNA expression of key inflammatory genes including *Tnf*, *Il6*, *Il12a*, *Il12b, Ifnb1* and *Cxcl9* was seen in *fe*s^-/-^ compared to *fes^+/+^* BMDMs (Fig.5e) and others (Extended Data Fig.5a-c). The differentially expressed genes used in a GO analysis were significantly associated with increased inflammatory pathway activation and cytokine production, including IL-12 binding and T cell activation (Extended Data Fig.5d). Exploring this further at the protein level, we confirmed higher secreted levels of 11 of the 13 cytokines seen to be transcriptionally upregulated in *fes^-/-^* compared to *fes^+/+^* BMDMs by RNAseq analysis (Fig. 5f-i; Extended Data Fig.5e-m); most notably TNFα, IL-6, IL-12, and IFNβ (Fig.5f-i).

Interestingly, *Il12b* was one of the of the most differentially expressed cytokines at 4 hours post-LPS stimulation (Fig.5e), and IL12 appeared in the GO analysis (Extended Data Fig.5d). However, secreted protein levels of IL-12p70 were very low at 4 hours, appeared at 8 hours, and showed a significant increase in *fes^-/-^* compared to *fes^+/+^* BMDMs at 12 hours (Fig.5h). One possible explanation of this is the potential for IL-12 to be secreted or remain membrane-bound; in which form it, has been shown to have improved CTL priming functionalities^37–39^. To explore the possibility that Fes might contribute to regulation of IL-12 trafficking, BMDMs were treated with LPS for 8hrs or 24hrs, then either permeabilized or left unpermeabilized to assess total or cell surface IL-12 or TNFα by flow cytometry. At 24 hours, there was an LPS-dependent increase in total IL-12 in both genotypes, but this was significantly higher in *fes^-/-^* BMDMs (Fig.5j). However, surface levels showed an LPS-induced increase in IL-12 only on *fes^-/-^*BMDMs (Fig.5k). This genotype difference in total or surface levels of IL-12 was not apparent at 8 hours post-LPS (Extended Data Fig.6a,b). A parallel analysis of TNFα, showed LPS-induced total levels at 8 hours, which were significantly higher in *fes^-/-^*BMDMs (Extended Data Fig.6c); and unlike IL-12 these total levels of TNFα had returned nearly to base line by 24 hours (Extended Data Fig.6e). In further contrast to IL-12, TNFα surface levels were not appreciably detected on either *fes^+/+^* or *fes^-/-^*BMDMs at either time point, regardless of LPS stimulation (Extended Data Fig.6d,f). These observations provide evidence for a novel cytokine-specific trafficking role for Fes.

### Fes deficient BMDMs displayed improved CTL priming, cancer cell toxicity, and tumour control

Increased TLR4 signaling and pro-inflammatory cytokine production by *fes^-/-^* BMDMs suggests improved CTL priming capability compared to *fes^+/+^*BMDMs. To explore this, we pulsed BMDMs with the chicken ovalbumin SIINFEKL peptide and stimulated them with either the TLR4 agonist LPS (1µg/mL; Fig.6a,b; Extended Data Fig.7a-c) or the TLR1/2 agonist Pam3Cys (0.1µg/mL; Fig.6c,d) before co-culturing with splenic OT-I CD8+ T cells, which express a T cell receptor specific for the SIINFEKL peptide^40^. *fes^-/-^*BMDMs activated a greater percentage of CD8^+^ T cells both with and without LPS (Fig.6a) or Pam3Cys stimulation (Fig.6c). *fes^-/-^* BMDMs also primed these CD8^+^ T cells to a greater extent than *fes^+/+^* BMDMs in the presence of LPS (Fig.6b) or Pam3Cys (Fig. 6d), as determined by IFNγ MFI. This effect was preserved across a range of SIINFEKL concentrations (Extended Data Fig.6a-c) and showed no priming in the absence of SIINFEKL peptide (Extended Data Fig.6e). There was no apparent *fes* genome associated differences in the ability of BMDMs to present the SIINFEKL peptide on MHCI (Extended Data Fig.6e)

**Fig. 6:**
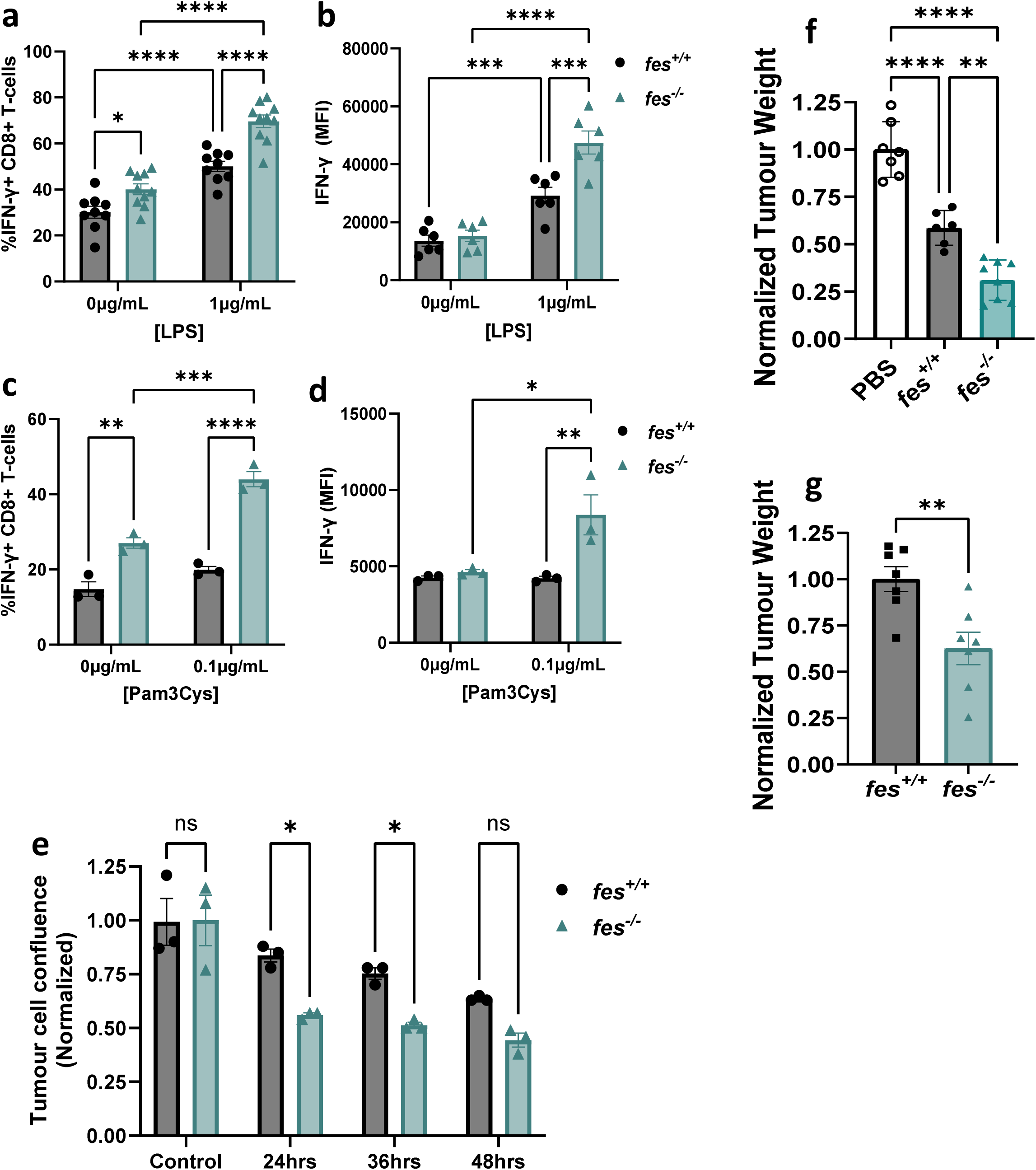
Fes deficient BMDMs displayed improved CTL priming, cancer cell toxicity, and tumour control. BMDMs were pulsed with 10^-10^M SIINFEKL peptide and stimulated with 1µg/mL LPS (**a**, *fes^+/+^* n=9 and *fes^-/-^* n=10; **b,** n=6 for each genotype); or 0.1µg/mL Pam3Cys (**c,** n=3 for each genotype; **d,** n=3 for each genotype) and co-cultured with OT-1 T cells. Cells were then analyzed for **(a,c)** intracellular IFNγ positivity and **(b,d)** IFNγ median fluorescence intensity (MFI). **(e)** BMDMs from either *fes^+/+^* or *fes^-/-^* mice (n=3) were co-cultured with GFP-expressing E0771 cells and either left untreated or treated with 1µg/mL LPS for 24, 36, or 48hrs. Confluency of the GFP positive E0771 cells was then calculated as fold change in confluency from 0hrs to the indicated time point and normalized to the untreated group. **(f)** 6e4 B16F10 cells or **(g)** 1e5 E0771 cells were orthotopically engrafted in *fes^+/+^* mice; and when tumours reached 100mm^3^, they were injected intratumourally with either **(f)** PBS, or **(f,g)** 3e6 *fes^+/+^* or *fes^-/-^* M1 polarized BMDMs. **(f)** Nine or **(g)** seven days later, mice were euthanized; and tumours weights measured and normalized to the **(f)** the PBS control group or **(g)** the *fes^+/+^* treated group. **a-d,f** were analyzed by two-way ANOVA with Tukey’s multiple comparisons test. **e** was analyzed by two-way ANOVA with Sidak’s multiple comparisons test, and **g** was analyzed by unpaired two-tailed t test; * p<0.05, ** p<0.01, *** p<0.001, **** p<0.0001.

Another important function of macrophages is the ability kill and phagocytose cancer cells to then promote adaptive immune responses^14,41,42^. To assess this, BMDMs were cultured with E0771 cancer cells for up to 48 hours in the presence or absence of LPS, which can promote macrophage cancer cell killing activity^14^. BMDMs had no differential genotype-associated effects on E0771 cell growth in the absence of LPS; but when LPS was added E0771 cells were prevented from proliferating to a greater degree in co-cultures with *fes^-/-^* BMDMs at 24 and 36 hours compared to 36 hours (Fig.6e).

Fes-deficiency in BMDMs enabled more efficient CTLs priming (Fig.6a-d), and conferred greater apparent suppression of cancer cell proliferation or survival (Fig.6e) suggesting Fes-deficiency in APCs may enable a reduction in tumour volume through both direct cancer cell killing and CTL priming. To directly show this, adoptive macrophage transfer experiments were performed with either *fes^+/+^* or *fes^-/-^* IFNγ-treated BMDMs injected directly into B16F10 (Fig.6f) or E0771 (Fig.6g) engrafted tumours in *fes^+/+^* mice. Both *fes^+/+^* and *fes^-/-^*BMDMs reduced B16F10 tumour weights compared to mock injections with PBS (Fig.6f); and *fes^-/-^* BMDMs were more effective at reducing tumour weights than *fes^+/+^* BMDMs in both models (Fig.6f,g).

## Discussion

We provide novel evidence that the Fes non-receptor tyrosine kinase plays an intracellular innate immune checkpoint role in APCs that modulates anti-cancer adaptive immunity in part by regulating expression and presentation of the pro-inflammatory Signal 3 cytokine IL-12 to CTLs. Using an *MMTV-Neu* transgenic breast cancer onset model we showed that targeted germline inactivating mutation in *fes* was associated with gene-dose dependent delay in tumour onset. This correlated with increased immune infiltration of CD3 and CD11b positive cells and inflammatory signaling (pNF-κB) prior to tumor detection. Inflammation can suppress tumour initiation and growth in some contexts (acute inflammation); while in others it can induce tumourigenesis (chronic inflammation)^43–45^. This is consistent with our understanding that the immune system can suppress tumourigenesis until the cancer overcomes this through induction of immunosuppression^46^. This was seen in *MMTV-Neu* mice, where following tumourigenesis, growth rates were identical regardless of *fes* genotype. This could potentially explain lower tumour incidence in people harbouring the rs17514846 SNP, which is associated with reduced Fes expression by ∼50%^29^.

*fes^-/-^* BMDMs displayed increased LPS-induced TLR4 signaling through NFκB, TBK1, and IRF3. This is important as IRF3 and NFκB are crucial for Signal 3 cytokine production^47^. RNAseq analysis indicated that Signal 3 cytokine gene transcription was more extensively upregulated in *fes^-/-^*versus *fes^+/+^* BMDMs. Transcripts encoding Signal 2 molecules such as CD80, TL1A and CD40^48–52^, were also more abundant in *fes^-/-^* BMDMs with and without LPS, potentially explaining the increased T cell activation by *fes^-/-^* BMDMs without LPS. Increased co-stimulatory proteins and enhanced sensitivity of *fes^-/-^* macrophages to LPS *in vitro*, and DAMPs *in vivo*, may have contributed to increased T cell activation in both E0771 and B16F10 models, including the ability of Dox-induced ICD to increase T cell activation. Tumour growth is associated with cancer cell death^53,54^, which may have produced sufficient levels of DAMPs to preferentially activate innate immunity in *fes^-/-^* mice. Fes is also expressed in vascular endothelial cells and its deletion has been previously correlated with attenuated tumour-associated angiogenesis^26,55^. In the current study, Fes deficiency in vascular endothelial cells may have contributed to slower tumour growth and increased necrosis due to lack of sufficient oxygen and nutrients^56^, as well as enhanced activation of Fes expressing innate immune cells in response to DAMPs released during necrosis. This is supported by the increased sensitivity of *fes^-/-^* BMDMs to low concentrations of LPS, and their enhanced *in vitro* CTL priming functionality at low antigen levels. This may also explain delayed tumor onset in the *MMTV-Neu* model, where *fes^KR/KR^*mice not only had more CD11b myeloid cells within their mammary glands prior to tumor onset, but *fes^KR/KR^* macrophages or DCs among these cells may have had increased anti-cancer APC functionality. Increased anti-tumor functionality of Fes-deficient BMDMs was also supported by adoptive macrophage transfer studies, where tumor-injected *fes^-/-^* BMDMs achieved greater tumor growth control than *fes^+/+^* BMDMs. Given the results of the *in vitro* BMDM/T cell co-culture priming experiments, this improved tumour growth control was likely due to increased *in vivo* inflammatory cytokine presentation and T cell priming capabilities.

While LPS-induced IL-12 expression at both the mRNA and secreted protein levels was higher in *fes^-/-^* than in *fes^+/+^* BMDMs, we were intrigued by the increased IL-12 surface retention phenotype of *fes^-/-^*BMDMs; which could contribute to improved T cell priming^37–39^. Increased membrane-associated IL-12 has been associated with improved DC-mediated priming of OT-II T cells, despite an apparent reduction in secreted IL-12^39^. This suggests that membrane-associated IL-12 may be more important than secreted IL-12 in stimulating T cells. This provides a potential novel mechanism by which Fes might regulate T cell priming; not only suppressing signaling from TLRs which drives IL-12 expression, but also by regulating post-translational trafficking, promoting more IL-12 secretion and less membrane retention, potentially leading to less effective T cell priming. Fes has been localized in the Golgi and within vesicles involved in both endocytic and exocytic pathways^25^. This is significant as IL-12 exists a membrane-bound protein until its cleavage in the Golgi^25,57,58^. In the absence of this cleavage step within the Golgi, IL-12 is retained on the cell surface^25,57,58^. IL-12 has also been shown to localize in late endosomes, and genetic disruption of VAMP7, a SNARE involved in exocytosis, caused a decrease in secreted IL-12^59^. Interestingly, Fes and its Fer paralog are the only two kinases containing an F-BAR domain, which shows conservation with CIP4 family adaptor proteins and selected RhoGAPs involved in membrane cytoskeletal dynamics, including receptor endocytosis^60,61^. Fes was implicated in regulating LPS-induced internalization of TLR4^21^, while Fer was shown to regulate ligand induced internalization of the EGFR^62^. These observations provide evidence supporting roles for Fes/Fer kinases in trafficking signaling proteins which are consistent with our observations regarding IL-12 in APCs. Future studies should explore molecular roles for Fes in IL-12 regulation within APCs, including potential effects on IL-12 post-translational modifications and trafficking.

The observed effect of Fes deficiency on sensitivity to anti-PD-1 therapy has important potential clinical implications. ICI therapy has become one of the most widely used therapies in cancer, with over 1500 clinical trials using anti-PD-1 antibodies. ICI therapy has been approved as first-line treatment in metastatic melanoma (32.6-45% response rate)^63–65^. It is also being used in other cancers, including TNBC in combination with chemotherapy (up to 21% response rate)^66–68^; and anti-PD-1 combined with chemotherapy improved the response rate by 13% over chemotherapy alone^69^. Another group showed that ICD induction with Dox, followed by anti-PD-1 treatment in TNBC achieved a response rate of 35%, but most interestingly, showed an increase in gene expression of immune-related genes involved in PD-1/PD-L1 and T cell cytotoxicity pathways^30^. This reprogramming of the tumour microenvironment to a more inflamed and anti-tumoural state was recapitulated in our study, which showed increased T cell activation and PD-1 expression in tumors in *fes^-/-^*, but not *fes^+/+^*mice. Furthermore, E0771 and B16F10 models are commonly reported in the literature as being resistant to anti-PD-1 therapy^31–36,70–72^. This was seen in our studies where even in combination with Dox, anti-PD-1 elicited only a marginal improvement on tumour growth, and no improvement on survival in *fes^+/+^* mice in either model. However, the combination of Dox and anti-PD-1 therapy in *fes^-/-^* mice conferred a greater delay in tumour growth and a significant increase in survival compared to Dox alone. This demonstrated that host Fes-deficiency sensitizes these tumours to anti-PD-1 therapy, most likely because of increased T cell activation and PD-1 positivity seen in *fes^-/-^*tumours at baseline and following Dox therapy. This is reminiscent of the aforementioned combined chemotherapy plus PD-1 study of metastatic TNBC which was reported in the TONIC clinical trial^30^. These results highlight Fes as both a potential therapeutic target as well as a predictive biomarker for successful ICI treatment.

Future studies should aim to elucidate Fes-mediated molecular mechanisms of IL-12 expression, trafficking, secretion, and presentation on the APC surface; correlate myeloid-specific Fes expression with patient outcomes through publicly available outcome data; and develop effective and specific inhibitors for Fes with limited off-target effects. The ALK inhibitor Lorlatinib, developed for use in non-small cell lung cancer, also inhibits Fes and several other kinases^73^; and it could be employed until more specific Fes inhibitors are available.

In our study, we have shown that genetic disruption of the non-receptor tyrosine kinase Fes enhances anti-cancer immunity, improving both chemotherapy and ICI immunotherapy efficacy. We show that Fes functions as an innate intracellular immune checkpoint, and its deletion or inactivation leads to increased anti-tumour immune cell tumour infiltration and activation. This change in immune cell composition of the tumour correlates with increased tumour control and survival in both TNBC and melanoma preclinical models, as well as sensitization to ICD-inducing chemotherapy and anti-PD-1 ICI therapy. These observations support a role for Fes as a tumour agnostic innate intracellular immune checkpoint and identify it as both a predictive biomarker for ICI immunotherapy success, and an attractive therapeutic target.

## Extended Data Fig. Captions

**Extended Data Fig.1: Tumour growth rates and representative IHC images, and pNFkB tumour immunoblotting.** *fes^+/+^*, *fes^+/KR^*, or *fes^KR/KR^* mice crossed with *MMTV-Neu* transgenic mice were monitored for tumour initiation by physical palpation, and tumours volumes were then assessed weekly by caliper measurements. **a,** Tumor growth rate constants were calculated as a function of changes volume per day. Representative images of immunohistochemical staining of mammary gland sections for; CD11b **(b)**, CD3 **(c)**, and Ki67 **(d)**. **e,** Immunoblotting analysis of pNFκB and Tubulin in lysates of premalignant glands from either *fes^+/+^*or *fes^-/-^ MMTV-Neu* transgenic mice. **(f)** Representative hematoxylin and eosin stained mammary gland section.

**Extended Data Fig.2: Fes is expressed in bone marrow derived macrophages, B cells, and NK cells, but not T cells. a,** Flow plots of negative isolation of T cells, NK cells, and B cells. **b,** Immunoblotting analysis of Fes expression in total splenocytes, BMDMs, negatively isolated T cells, negatively isolated B cells, negatively isolated NK cells, E0771 cells and B16F10 cells.

**Extended Data Fig.3: Representative immune profiling flow plots and splenic and tumour immune profiling.** Representative flow cytometry plots for immune profiling of **(a)** CD8^+^ T cells or **(b)** macrophage and dendritic cells. Spleens from the EO771 model **(c-j)** or B16F10 **(m-t)** were dissociated, stained and analyzed by flow cytometry for: **(c,m**) CD8+ T cells (CD45+, CD3+, CD8+ [CTLs]); **(d,n)** CD69+ CTLs; **(e,o)** PD-1+ CTLs; **(f,p)** macrophages (CD45+, CD11b+, F4/80+); **(g,q)** M2 macrophages (CD206+); **(h,r)** M1 macrophages (CD206-, MHCII+ and/or CD80+); **(i,s)** dendritic cells (CD45+, F480-, CD11c+, MHCII+); and **(j,t)** activated dendritic cells (CD80+). Tumours from the EO771 model **(k,l)** or B16F10 **(u,v)** were similarly analyzed for: **(k,u)** dendritic cells (CD45+, F480-, CD11c+, MHCII+); and **(l,v)** activated dendritic cells. Results were analyzed by one-way ANOVA with Tukey’s multiple comparisons test; * p<0.05, ** p<0.01, *** p<0.001, **** p<0.0001.

**Extended Data Fig.4: Fes-deficiency improves dendritic cell TLR signaling and increases sensitivity to LPS stimulation.** Bone marrow derived macrophages (BMDMs) were stimulated with the indicated concentrations of LPS for 120 minutes, and probed by immunoblot for (**a**) phospho-NFκB, and (**b**) phospho-TBK1 with densitometry normalized to the 27ng/mL well of *fes^+/+^* BMDMs.

**Extended Data Fig.5: Fes-deficiency confers increased cytokine expression. a-d,** BMDMs from either *fes^+/+^ (*n=3) or *fes^-/-^* mice (n=3) were untreated or stimulated with 1µg/mL LPS for 4 hours before RNA was isolated and subjected to poly-A enriched RNA sequencing analysis. The differential transcript levels of select interleukins **(a)**, C-C chemokine ligands **(b)**, and C-X-C chemokine ligands **(c)** were plotted. All genes that met the following cutoffs were then used for GO analysis **(d)** ^74^: (1) were LPS inducible; (2) had at least a 1.25 fold increase in gene expression in LPS stimulated *fes^-/-^* compared to *fes^+/+^* BMDMs; and (3) the fold change associated with LPS stimulation was greater in *fes^-/-^*compared to *fes^+/+^*BMDMs. **e-m,** BMDMs of either genotype were stimulated with 1µg/mL LPS for the indicated times and supernatants were assessed using the BioLegend Mouse Inflammation LEGENDplex panel. Selected analytes displaying differential expression included: CCL2 **(e)**, GM-CSF **(f)**, IFNγ **(g)**, IL-1α **(h)**, IL-1β **(i)**, IL-10 **(j)**, IL-17A **(k)**, IL-23 **(l)**, and IL-27 **(m)**. Results were analyzed by two-way ANOVA with Sidak’s multiple comparisons test; * p<0.05, ** p<0.01, *** p<0.001, **** p<0.0001.

**Extended Data Fig.6: Fes-deficient BMDMs show time-dependent changes in cytokine expression and localization. a,b,** BMDMs isolated from either genotype (n=3) were stimulated for 8 hours with 1µg/mL LPS and assessed by flow cytometry for either **(a)** total IL-12 (staining after permeabilization), or **(b)** cell surface IL-12 (staining prior to permeabilization), respectively. **c-f**, TNFα was similarly assessed at **(c,d)** 8 hours (n=6) and **(e,f)** 24 hours (n=3). Results were analyzed by two-way ANOVA with Tukey’s multiple comparisons test; * p<0.05, ** p<0.01, *** p<0.001, **** p<0.0001.

**Extended Data Fig 7: Fes-deficiency improves T cell priming capabilities of BMDMs at a range of SIINFEKL concentrations in a Signal 2-independent mechanism.** The relative ability of BMDMs from either *fes^+/+^*(n=9) or *fes^-/-^* (n=9) mice to activate OT-I T cells was assessed after pulsing the BMDMs with **(a)** 10^-9^M, **(b)** 10^-11^M, **(c)** 10^-12^M, or **(d)** no SIINFEKL peptide. **e,** the relative ability of *fes^+/+^* (n=5) or *fes^-/-^* (n=5) BMDMs to present SIINFEKL on MHCI was assessed by flow cytometry after pulsing with 10^-10^M SIINFEKL for 2 hours, washing, and stimulating or not with 1µg/mL LPS for 7 hours and staining with a H-2Kb bound to SIINFEKL antibody. Results were analyzed by two-way ANOVA with Tukey’s multiple comparisons test; * p<0.05, ** p<0.01, *** p<0.001, **** p<0.0001.

## Methods

This research complies with all relevant institutional ethical and biosafety regulations. Mouse experiments were approved by the Queen’s University Animal Care Committee and conform to the Canadian Council on Animal Care guidelines.

### Cell lines

E0771 (CRL-3461), 293T (CRL-3216) and B16F10 (CRL-6475) were purchased from the American Type Culture Collection (ATCC, Manassas, VA). E0771, B16F10 and 293T cells were cultured in Dulbecco’s Modified Eagle Medium (DMEM; Gibco cat#: 12100-061) supplemented with 10% fetal bovine serum (FBS; Corning Cat#:35-077-CV) 2mM L-glutamine (Corning; cat#:25-005-CI), and 1% antibiotic and antimycotics (A/A. Gibco; cat#:15240-062). Cell lines were cultured in a humidified incubator held at 5% CO_2_ and 37℃. Cells used in all experiments were determined to be mycoplasma negative by the mycoplasma PCR detection kit (Applied Biological Materials; cat# G238).

### Generation of bone marrow derived macrophages and dendritic cells

Bone marrow derived macrophages (BMDMs) were isolated as described previously^75,76^ Briefly, on day 1, *fes^+/+^* or *fes^-/-^* ^20^C57BL/6 mice (8 – 12 weeks old) were sacrificed and bone marrow flushed from both tibias and femurs into 5mL of Roswell Park Memorial Institute (RPMI) 1640 (Sigma-Aldrich; cat#:R8758-500mL) supplemented with 2% FBS and 1% A/A. Isolated bone marrow was dispersed by passing through a 23-gauge needle. Cells were then passed through a 40µm cell strainer and pelleted at 1000rpm for 5 minutes. Cells were then resuspended in 10mL BMDM media (RPMI 1640 supplemented with 15% FBS, 1% A/A, 50µM α-monothioglycerate [α-MTG; Sigma; cat#:M-1753], and 20ng/mL macrophage colony-stimulating factor [M-CSF; a generous gift from Richard Stanley]^77^) and plated in 100mm cell culture plates. On day 2, non-adherent cells were collected and pelleted at 1000rpm for 5 minutes. Cells were resuspended in 6mL of BMDM media (3mL fresh media and 3mL of conditioned media from previous day) and plated on a 60mm cell culture plate. On Day 4, non-adherent cells were collected, pelleted, resuspended in 10mL of BMDM media, and plated onto untreated tissue culture plates. After 72 hours, cells were trypsinized, counted, and used for experiments.

### Immunoblotting analysis of LPS signaling

BMDMs from *fes^+/+^* and *fes^-/-^* mice were plated in 6-well flat bottom TC-treated plates (Corning, USA) at 2,500,000 cells per well in 2mL of complete macrophage media. The BMDMs were either left untreated or were stimulated with 1μg/mL Lipopolysaccharides (LPS; *E. coli*, serotype 055:B5, Sigma Aldrich cat#: L2637), for 15, 30, 45, or 60 minutes. Cells were washed twice with ice-cold phosphate buffered saline (PBS; 170mM NaCl, 3.5mM KCl, 10mM Na_2_HPO_4_, 2mM KH_2_PO_4_, pH 7.2) supplemented with 2mM sodium orthovanadate (ThermoFisher Scientific cat#: 205330500) and lysed in RIPA buffer (10 mM Tris pH7.2, 158 mM NaCl, 1 mM EDTA, 0.1% SDS, 1% Na Deoxycholate, 1% Triton-100) supplemented with Halt^TM^ Protease Inhibitor Cocktail (ThermoFisher Scientific cat#: 87785) and sodium orthovanadate, or in 2x SDS Laemmli Sample Buffer for pIRF3 analysis. Protein quantification was performed using the Pierce^TM^ BCA Protein Assay Kit (ThermoFisher Scientific cat#: 23225) and immunoblotting was performed as previously described^78^ using the following antibodies and concentrations: Fes (1:1000; Cell Signaling Technologies cat#: 85704S), phospho-NF-κB (1:1000; Cell Signaling Technologies cat#: 3033S),, phospho-TBK1 (1:1000; Cell Signaling Technologies cat#: 5483),), phospho-IRF3 (1:1000; Cell Signaling Technologies cat#:29047S), γ-tubulin (1:1000; Sigma-Millipore, cat#: T5326), and GAPDH (1:10000; Cell Signaling Technologies cat#: 5174S).

### RNA sequencing

BMDMs from *fes^+/+^* and *fes^-/-^* mice were plated in 6-well flat bottom TC-treated plates (Corning, USA) at 2,500,000 cells per well in 2mL of complete macrophage media. The BMDMs were either left untreated, or they were stimulated with 1μg/mL LPS (Sigma-Aldrich, USA) for 4 hours. Cells were then lysed with 350μL of cold TRI Reagent® (Sigma-Aldrich, USA) for five minutes. RNA purification was performed according to Invitrogen’s TrizolTM Regent protocol. Purified RNA sample concentrations were determined with a NanoDrop ND-1000 spectrophotometer for downstream cDNA synthesis. cDNA synthesis was carried out using the Maxima First Strand cDNA synthesis kit (Thermo Fisher Scientific, USA) according to manufacturer’s instructions, with 372ng of RNA as the starting template. Samples were mixed, centrifuged, and placed in an Applied Biosystems thermocycler (Life Technologies, USA) for one cycle for 10 minutes at 25°C, 15 minutes at 50°C and finally 5 minutes at 85°C. RNA quality and quantity were confirmed via NanoDrop measurement. All samples possessed an A260/280 purity ratio >1.7 and were sent to Genome Quebec (Montreal, Canada) for poly-A enriched RNA-sequencing. Genome Quebec conducted an alignment of the sequencing data to the GRCm38 (Ensembl 102 release) murine genome to determine raw expression gene counts and transcripts per million (TPM) counts. The data was then pre-processed by removing genes with zero counts across all samples and filtering out genes with expression levels below the 75th percentile threshold (roughly 12 counts). The resulting pre-processed TPM counts for each gene were plotted for each genotype condition.

### LEGENDplex cytokine assay

BMDMs from *fes^+/+^* and *fes^-/-^* mice were plated in 6-well flat bottom TC-treated plates (Corning, USA) at 2,500,000 cells per well in 2mL. BMDMs were either left untreated, or they were stimulated with 1μg/mL LPS (Sigma-Aldrich, USA) in 1mL for 4 hours, 8 hours, or 12 hours. Supernatant was then collected, and spun at 10,000rpm for 12 minutes at 4°C to pellet cell debris. Supernatant was then used in the Mouse Inflammation LEGENDplex panel (BioLegend; cat #: 740446) according to manufacturer’s instructions.

### Macrophage and T cell co-culture

BMDMs were harvested as described above and were seeded in a 96-well plate. BMDMs were then treated for 2 hours with varying concentrations of OVA peptide (SIINFEKL: CPC Scientific cat#: MISC-012). BMDMs were then washed three times with serum free RPMI, before treatment with 1µg/mL of LPS (Sigma-Aldrich, USA) for 1 hour. BMDMs were then washed three times with serum free RPMI. Spleens from OT-I mice were isolated, manually dissociated, filtered through a 40µm filter, and subjected to CD8^+^ T cell negative selection with the EasySep^TM^ Mouse T cell isolation kit (STEMCELL Technologies cat#: 19853), according to the manufacturer’s protocol. Isolated CD8^+^ T cells were seeded onto treated BMDMs in a ratio of 1:1 resuspended in T cell media (RPMI 1640, 10% FBS, 1% A/A, 1% gentamicin (ThermoFisher Scientific cat#: 15710064) and incubated for 2 hours; after which cells were treated with 5μg/mL Brefeldin A (BioLegend; cat#: 420601) and incubated for a final 4 hours before recovery and analysis by flow cytometry.

### Animal Studies

All experiments involving the use of animals were performed in accordance with the Canadian Council on Animal Care guidelines and with the approval of Queen’s University Animal Care Committee. All mouse strains (*fes^KR/K^)*^28^, (*fes^-/-^*)^20^ and OT-I (C57BL/6-Tg(TcraTcrb)1100Mjb/J, https://www.jax.org/strain/003831)^40^ were on a C57BL/6 background. Mice were bred in-house and 8–12-week-old males or females were used in engraftment studies for B16F10 or E0771 models, respectively, with drug and antibody doses normalized to body weights. Animals were randomized at treatment start time, with any mice lacking tumour growth excluded from the study. Tumour length (*L*) and width (*W*) were measured by caliper, and tumour volume (*V*) calculated as *V* = (*L* x *W*^2^)/2. Tumours were measured until an endpoint of 1.1cm^3^ or humane endpoints (e.g., ulceration) at which point, mice were euthanized and survival analyzed.

### Engraftment models

C57BL/6 mice (8-12 weeks old) were orthotopically engrafted with 1×10^5^ E0771 cells resuspended in 20µL of a 1:1 ratio of Matrigel (Corning; cat#: 354234) to PBS injected into the fourth mammary fat pad with a Hamilton syringe; or with 6×10^4^ B16F10 cells resuspended in 50µL of a 1:1 ratio of Matrigel to PBS cell suspension injected subcutaneously with a 26-gauge needle into the left flank. Once tumours reached ∼100mm^3^ (E0771 model: *fes^+/+^* ∼12 days; *fes^-/-^* ∼14 days; B16F10 model: *fes^+/+^* ∼8 days; *fes^-/-^* ∼10 days), E0771 tumour-bearing mice were treated with either 5mg/kg of Doxorubicin (Dox: Selleckchem cat#: S120813) or vehicle control in a 200µL 5% DMSO/PBS solution injected intraperitoneally (IP), repeated 5 and 10 days later; or for the B16F10 model, 2.5mg/kg Dox or vehicle control in a 200µL 5% DMSO/PBS solution injected IP every 2 days to a final cumulative dose of 20mg/kg. Tumour growth and survival were analyzed as described above.

### Anti-PD-1 treatment

The anti-PD-1 treatment model followed a similar protocol to that described above; however, every mouse was treated with its tumour-specific Dox dose in combination with either 10mg/kg anti-IgG control antibody (BioXCell; cat#: BE0089) or 10mg/kg anti-PD-1 antibody (BioXCell; cat#:BE0146) IP, for a total of three doses, either with every Dox treatment in the E0771 model or every two Dox treatments starting on the third Dox treatment in the B16F10 model.

### Harvesting tumours and spleens for tumour immune profilin**g**

Mice were engrafted with cancer cells and treated as described above. However, three days following the second dose of Dox treatment in the E0771 model, or one day after the fourth dose in the B16F10 model, mice were euthanized, and tumours and spleens harvested. Tumours were cut into ∼1mm^3^ chunks with a straight razor, while spleens were pressed through a sieve with a syringe plunger. Tumour samples were suspended in RPMI 1640 supplemented with 10% FBS, 1mg/mL collagenase B (Sigma-Aldrich cat#: 11088815001) and 25µg/mL of DNAse I (Sigma-Aldrich Cat#: 10104159001); while splenocytes were resuspended in RPMI containing only 25µg/mL of DNAse I. Tumour chunks were then transferred into C Tubes (Miltenyi Biotec cat#: 130-096-334) and dissociated using the gentleMACS Dissociator following the manufacturer’s protocol for tough (E0771) or soft (B16F10) tumours. Tumour and spleen cell suspensions were then filtered through 70µM cell strainers, followed by 40µM cell strainers, and then treated with Ammonium Chloride Potassium (ACK) lysis buffer to lyse red blood cells within the preparation. Cells were then washed and resuspended in fluorescence-activated cell sorting (FACS) buffer (PBS containing 0.75% bovine serum albumin [BSA; BioShop cat#: ALB001], 5mM EDTA in PBS) in preparation for staining and flow cytometry.

### Antibody staining and flow cytometry

Harvested cells were plated into 96-well round bottom plates (1×10^5^ cells/well) and centrifuged at 1000rpm for 5 minutes. Supernatants were carefully decanted, and cell pellets resuspended in 100µL FACS containing 0.2µg TruStain FcX™ (anti-mouse CD16/32) antibody or 100µL of PBS containing 0.2µg TruStain FcX™ and 0.04µL Zombie AquaTM Fixable Viability Kit (BioLegend cat#: 423102) and incubated at room temperature in the dark for 30 minutes. Next, 200µL of FACS buffer was added to each well, gently mixed, and pelleted at 1000rpm for 5 minutes, and carefully decanted. Next, cells were stained by resuspending in 100mL of FACS buffer containing appropriate antibody panels (listed in Supplementary Table 1) and incubated on ice in the dark for 30 minutes. 200µL of FACS was added to each well, gently mixed, and pelleted at 1000rpm for 5 minutes, and carefully decanted, followed by an additional wash in 200µL FACS buffer. If the cells were not being subjected to intracellular staining, cells were resuspended in 100µL of 1% paraformaldehyde in PBS and incubated at room temperature for 30 minutes. An additional 200µL of FACS buffer was added, cells pelleted at 1500rpm for 5 minutes, decanted, and resuspended in 200µL FACS buffer to be analyzed on a CytoFlex-S Flow Cytometer (Beckman Coulter). For intracellular staining, cells were stained following the ThermoFisher “BestProtocols: Staining Intracellular Antigens for Flow Cytometry – Protocol A” using the eBioscience^TM^ Permeabilization Buffer (ThermoFisher Scientific cat#: 00-8333-56) and eBioscience^TM^ IC Fixation Buffer (ThermoFisher Scientific cat#: 00-8222-49) before analysis on the CytoFlex-S Flow Cytometer (Beckman Coulter). Data analysis was performed using FlowJo® v10 software (FlowJo, LLC, Ashland, OR).

### Macrophage adoptive transfer

C57BL/6 *fes^+/+^* mice (8-12 weeks old) were engrafted with either E0771 or B16F10 cells as described above. BMDMs for adoptive transfer were isolated from 8-12 weeks old *fes^+/+^*or *fes^-/-^* mice as described above. On day 6, the BMDMs were treated with 20ng/mL interferon gamma (IFN-γ; Peprotech cat#: 315-05) and left overnight to polarize them into an M1-like state by Day 7. On Day 7, BMDMs were harvested using Cellstriper (Corning; cat#: 25-056-CI), washed with PBS and 3×10^6^ *fes^+/+^* or *fes^-/-^* M1-polarized BMDMs resuspended in 50μL of PBS, or PBS alone, were injected intratumourally (IT) with a 26-gauge needle into 100mm^3^ tumours. Seven or nine days later (for EO771 or B16F10 tumors, respectively), mice were euthanized, and tumour weights determined.

**Supplementary Table 1:** Immune profiling antibody panels.

| T cell panel |  |  |  |
| --- | --- | --- | --- |
| Target | Fluorophore | Vendor | Catalog # |
| Viability | Aqua (Krome Orange) | Invitrogen | L34966 |
| CD45 | FITC | Biolegend | 103018 |
| CD3 | Alexa Fluo® 700 | Biolegend | 100216 |
| CD8 | PE | Biolegend | 100708 |
| CD69 | PE/Cy5 | Invitrogen | 15-0691-82 |
| PD-1 | APC/Cy7 | Biolegend | 135224 |

| Macrophage and Dendritic cell panel |  |  |  |
| --- | --- | --- | --- |
| Target | Fluorophore | Vendor | Catalog # |
| Viability | Aqua (Krome Orange) | Invitrogen | L34966 |
| CD45 | FITC | Biolegend | 103018 |
| CD11b | Alexa Fluo® 700 | Biolegend | 101222 |
| F4/80 | PE/Cy7 | Biolegend | 123114 |
| MHC-II | Pacific Blue 450 | Biolegend | 107620 |
| CD11c | BV605 | Biolegend | 117333 |
| CD80 | PE | Biolegend | 104708 |
| CD206 | PE/Dazzle 594 | Biolegend | 141732 |
| PD-1 | APC/Cy7 | Biolegend | 135224 |
| Additional Antibodies |  |  |  |
| Target | Fluorophore | Vendor | Catalog # |
| CD8 | FITC | Biolegend | 100706 |
| Interferon $\gamma$ | PE | Biolegend | 505808 |
| F4/80 | APC/Cy7 | Biolegend | 123118 |
| Interleukin-12 p35 | PE | Invitrogen | MA5-23559 |
| TNF-α | Alexa Fluor® 488 | Biolegend | 506313 |
| H-2K<sup>b</sup> bound to SIINFEKL | APC | Biolegend | 141606 |

