## Supplemental Figures for "Fes-deficient macrophages enhance CD8^+^ T cell priming and tumour control through increased pro-inflammatory cytokine production and localization"

### Slide 1
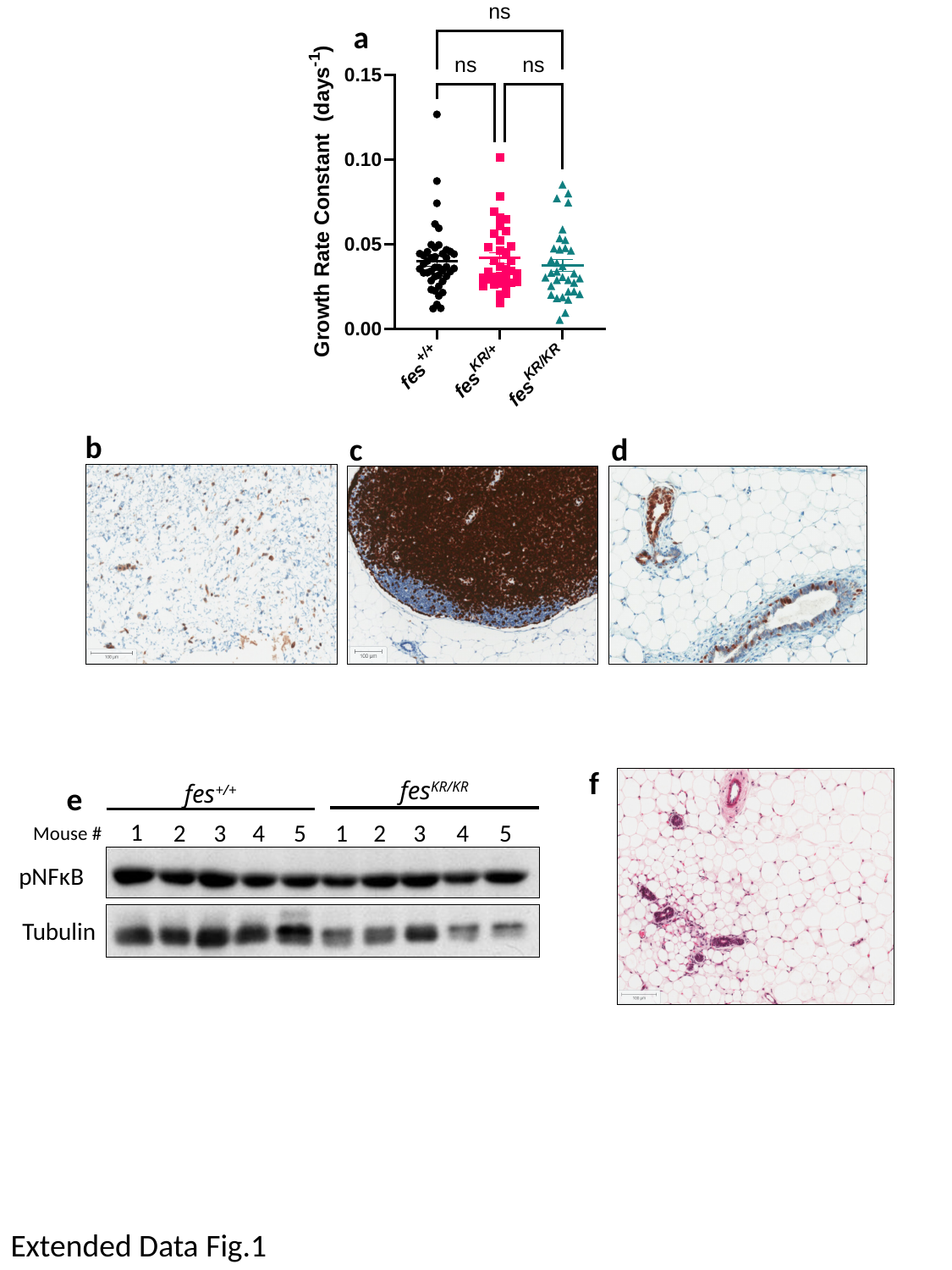

a
b
c
d
f
fes^KR/KR^
fes^+/+^
e
1
4
2
5
4
3
2
5
1
3
Mouse #
pNFκB
Tubulin
Extended Data Fig.1

### Slide 2
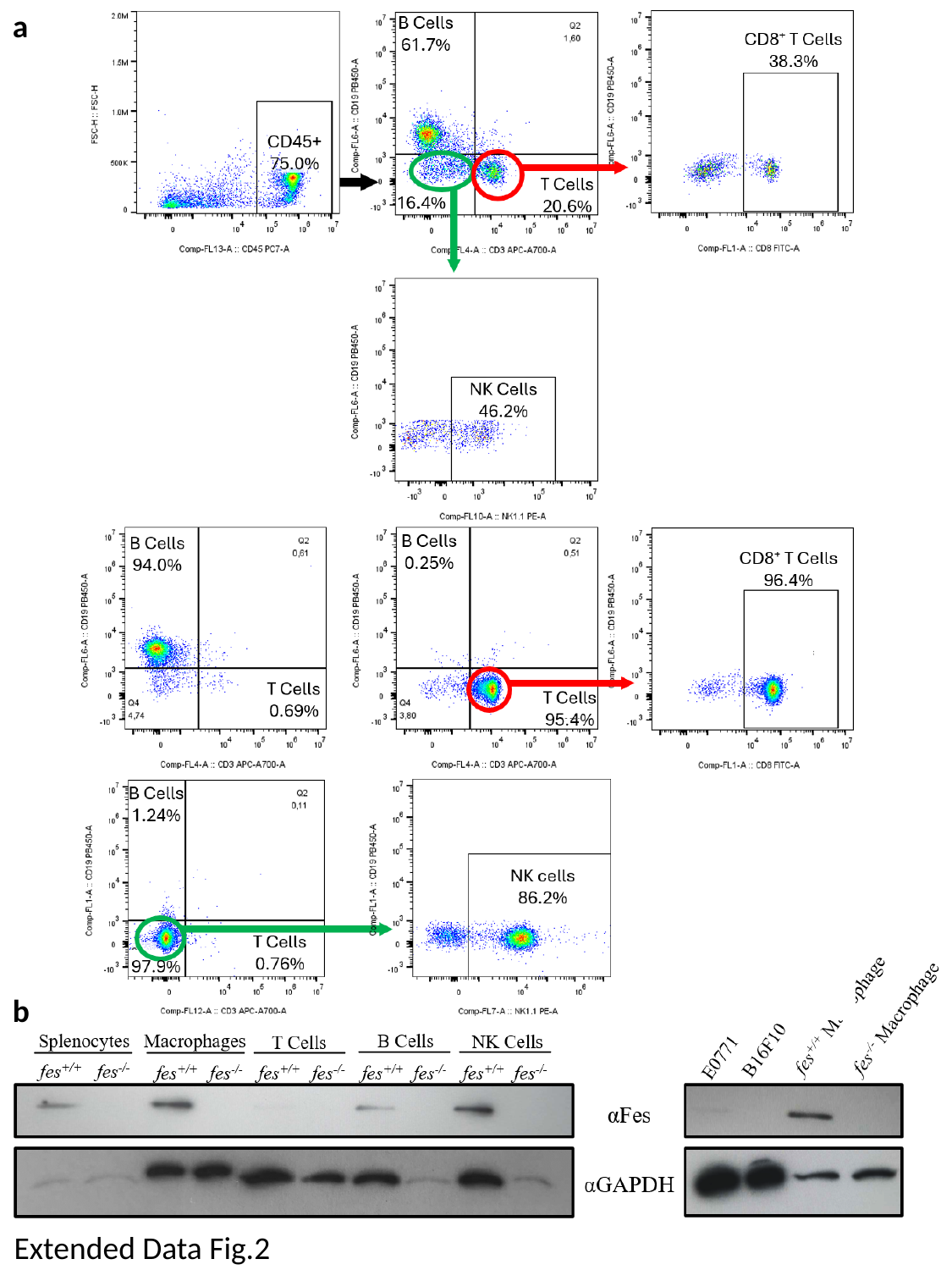

a
b
Extended Data Fig.2

### Slide 3
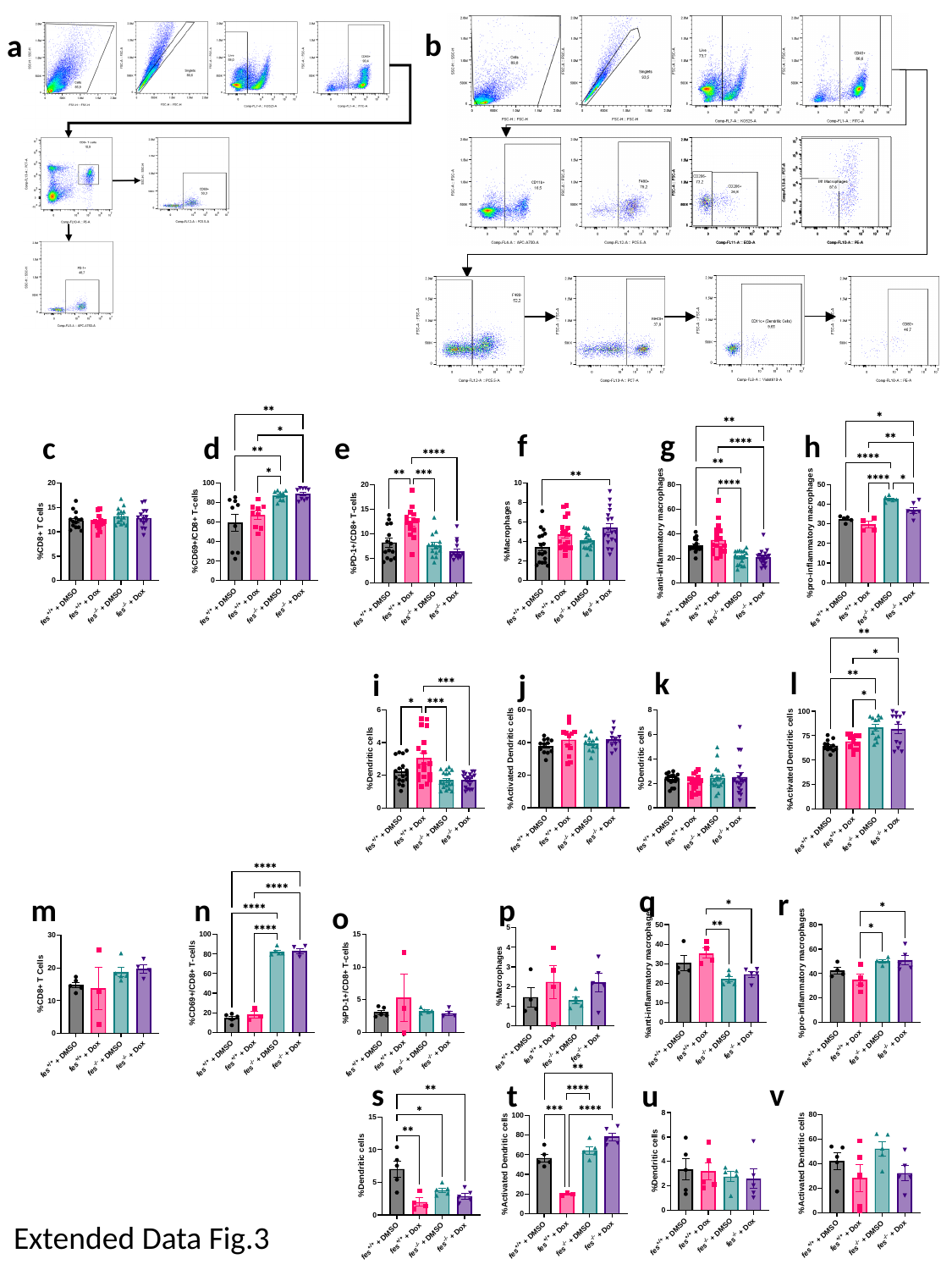

b
a
g
f
h
c
d
e
k
l
i
j
q
r
p
m
n
o
s
v
u
t
Extended Data Fig.3

### Slide 4
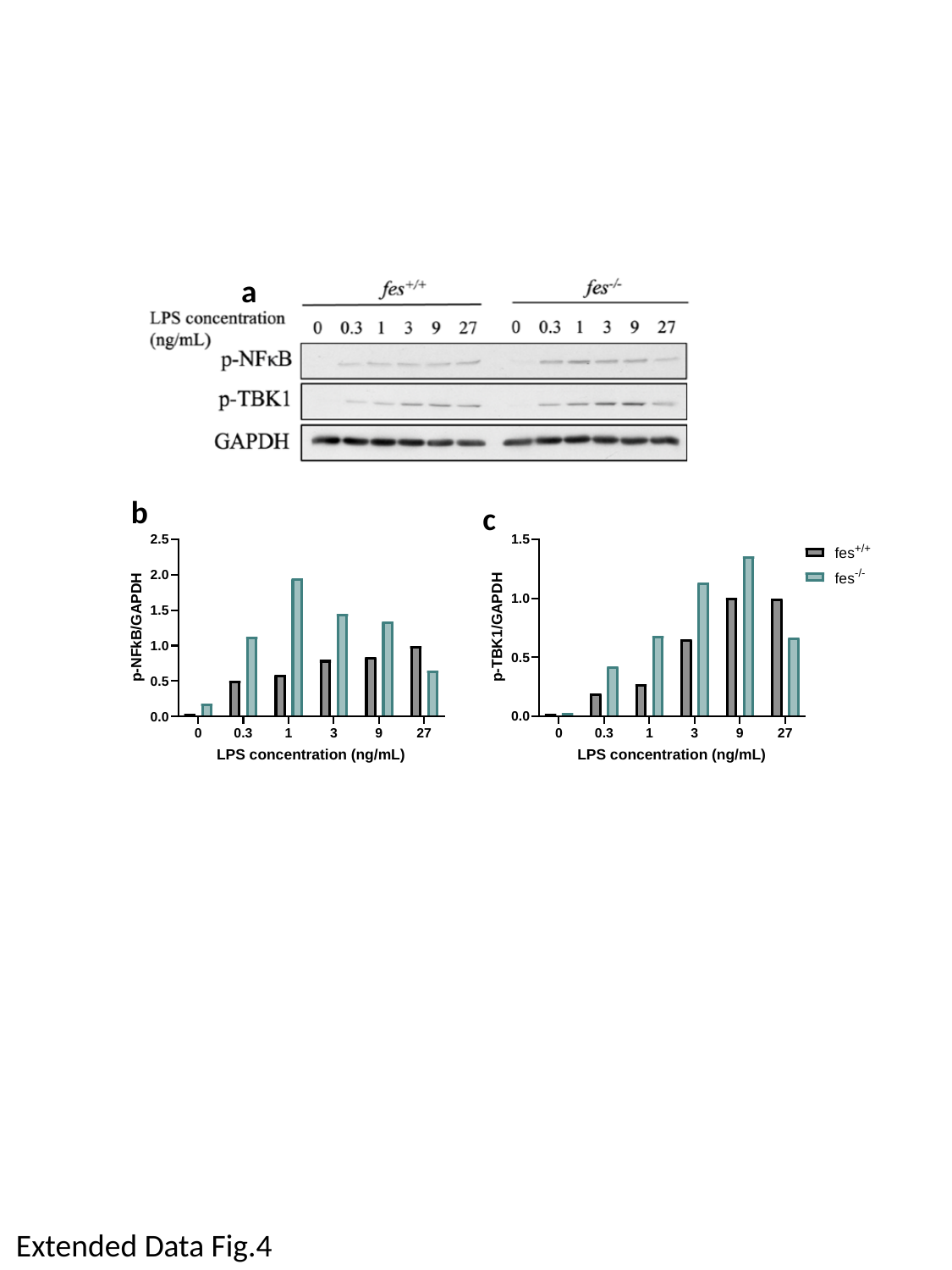

a
b
c
Extended Data Fig.4

### Slide 5
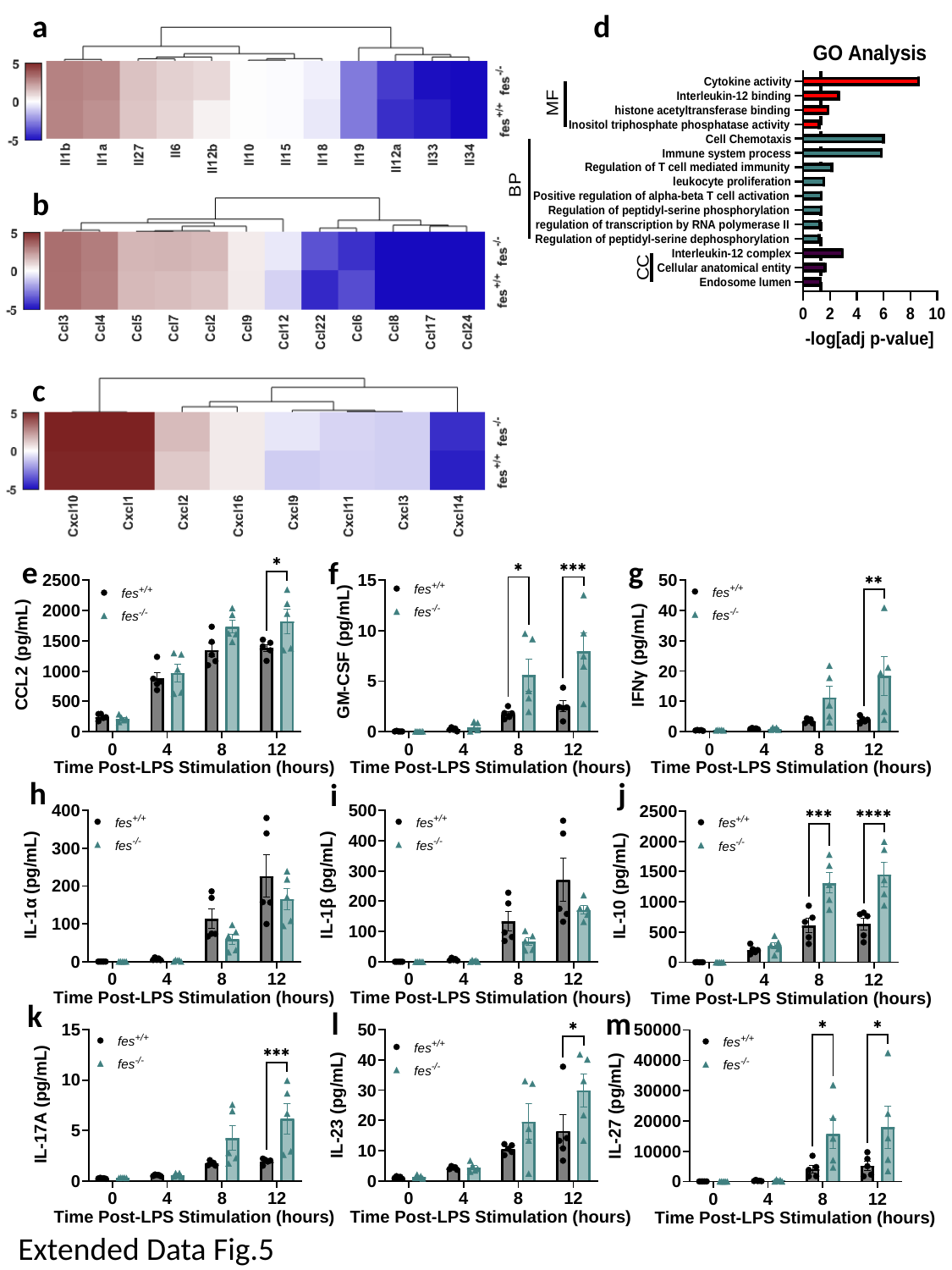

a
d
b
c
e
g
f
h
j
i
k
l
m
Extended Data Fig.5

### Slide 6
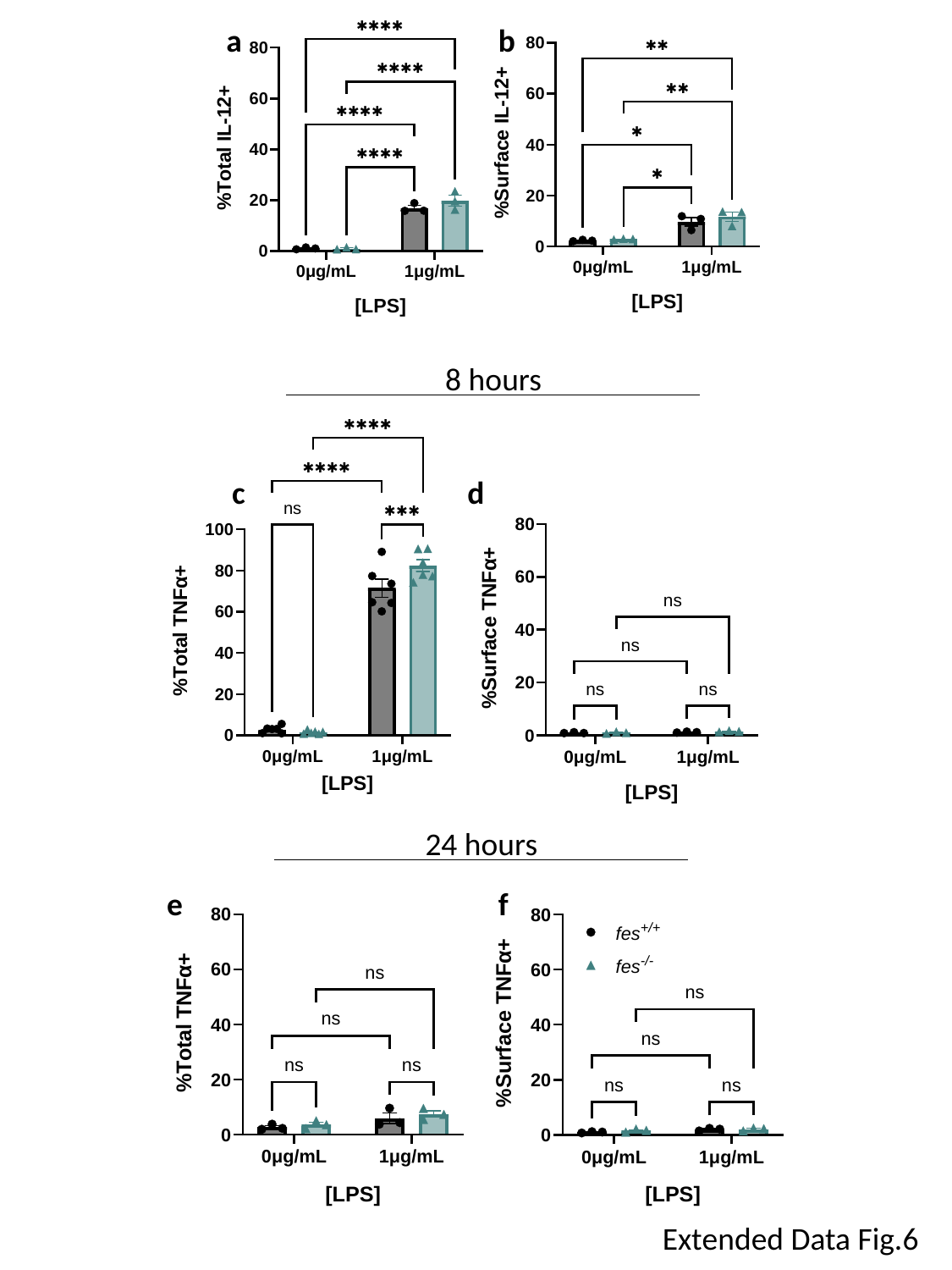

a
b
8 hours
c
d
24 hours
e
f
Extended Data Fig.6

### Slide 7
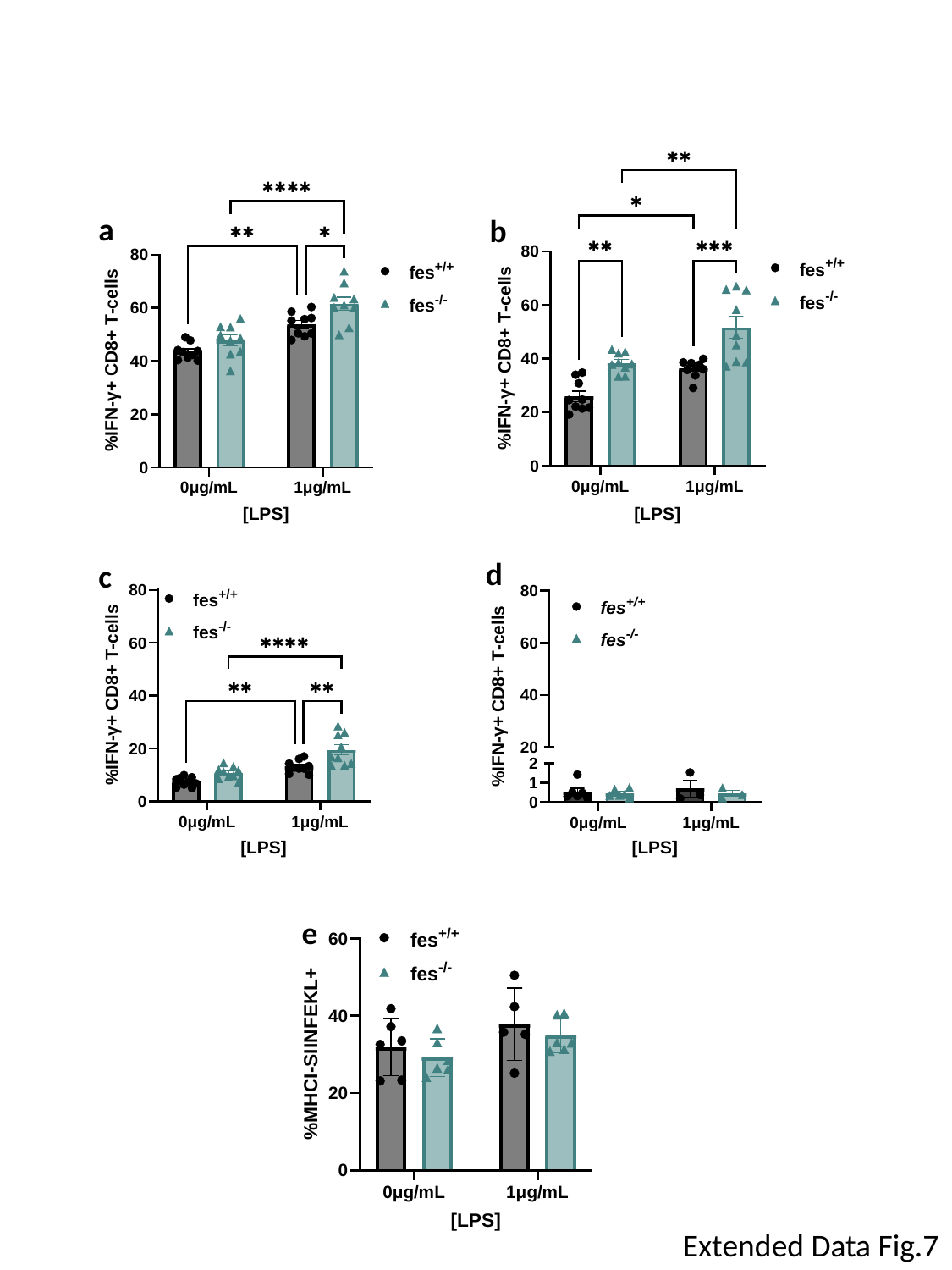

a
b
d
c
e
Extended Data Fig.7
